# Structure of the peroxisomal Pex1/Pex6 ATPase complex bound to a substrate

**DOI:** 10.1101/2022.11.19.517173

**Authors:** Maximilian Rüttermann, Michelle Koci, Pascal Lill, Björn Udo Klink, Ralf Erdmann, Christos Gatsogiannis

## Abstract

The double-ring AAA+ ATPase Pex1/Pex6 is required for peroxisomal receptor recycling and is essential for peroxisome formation. Pex1/Pex6 mutations cause severe peroxisome associated developmental disorders. Despite its pathophysiological importance, mechanistic details of the heterohexamer are not yet available. Here, we report cryoEM structures of Pex1/Pex6 from *Saccharomyces cerevisiae*, with an endogenous protein substrate trapped in the central pore of the second ring (D2). Pairs of Pex1/Pex6(D2) subdomains engage the substrate via a staircase of pore-1 loops with distinct properties. The first ring (D1) is catalytically inactive but undergoes significant conformational changes resulting in alternate widening and narrowing of its pore. These events are fueled by ATP hydrolysis in the D2 ring and disengagement of a “twin-seam” Pex1/Pex6(D2) heterodimer from the staircase. Mechanical forces are propagated in a unique manner along Pex1/Pex6 interfaces that are not available in homo-oligomeric AAA-ATPases. Our structural analysis reveals the mechanisms of how Pex1 and Pex6 coordinate to achieve substrate translocation.

## Introduction

Peroxisomes are single-membrane enclosed eukaryotic organelles playing central roles in lipid metabolism and maintenance of redox balance^1–3^. Impaired peroxisomal assembly and/or defects in reactive pathways housed by peroxisomes manifest in severe metabolic disorders^4–6^. There is also increasing evidence correlating peroxisomal dysfunction with aging and a broad range of relevant diseases, including cancer, Parkinson’s disease and diabetes^7–9^. Peroxisomal enzymes are produced in the cytosol and then imported into the peroxisomal matrix^10^. The majority of peroxisomal matrix enzymes carry a C-terminal <u>P</u>eroxisome <u>T</u>argeting <u>S</u>ignal (PTS1) motif, which is recognized by the peroxisomal receptor Pex5 in the cytosol^11^. Pex5 delivers its cargo to a docking complex at the peroxisomal membrane^12,13^. This interaction triggers the formation of a transient channel for cargo translocation along the peroxisomal membrane^14,15^. The underlying mechanism is not yet understood.

It is thought that during this process, the receptor either enters the membrane to become part of the channel (similar to a pore-forming toxin) or accompanies the cargo and enters completely into the peroxisomal lumen^15–17^. For the next round of import, the receptor has to be recycled and thus translocated back to the cytosol^16,18^. The peroxisomal ubiquitin-ligase complex Pex2/Pex10/Pex12 was recently shown to contain a pore that might provide the retro-translocation path for the receptor^19^. The unstructured N-terminal peptide of the receptor was suggested to enter this pore from the peroxisomal lumen, and subsequently get mono-ubiquitylated by the ring-finger peroxin Pex2 of the ubiquitin ligase complex^19,20^. The Pex5 ubiquitin moiety is subsequently recognized by the type II (AAA)+ heterohexameric complex Pex1/Pex6, which is attached to the peroxisomal membrane via the tail anchored protein Pex15 (Pex26 in mammals)^21–23^. Under ATP consumption, Pex1/Pex6 is expected to process Pex5 by pulling it through its central ATPase pore back to the cytosol^23,24^. Receptor recycling by Pex1/Pex6 consumes energy, but Pex5 import is ATP independent^25,26^. However, both machineries (cargo import and receptor export) are functionally linked via Pex15^27^ and the ubiquitination of Pex5^18,28^. This supports the idea of an export-driven peroxisomal enzyme import, with Pex1/Pex6 being the driving molecular motor for a sustainable import^29–31^. When Pex1 and/or Pex6 are impaired, ubiquitinated Pex5 accumulates at the peroxisomal membrane, which triggers pexophagy^20,32^. More recently, the autophagy receptor Atg36 has been proposed as a novel Pex1/Pex6 substrate in yeast^32^. This suggests that Pex1/Pex6 might have additional functions in organelle quality control^33^, which may explain the mostly lethal phenotype of peroxisomal biogenesis disorders (PBDs)^33^.

Pex1 and Pex6 are both composed of two N-terminal domains (N1 and N2) followed by two AAA cassettes, known as the D1 and D2 domains (Fig. 1a). Each cassette is comprised of two distinct subdomains: a core nucleotide-binding domain (large ATPase subdomain, _L_D1/ _L_D2) and a smaller domain of α-helical bundles (small ATPase domain, _S_D1/ _S_D2). The tandem N-terminal domains (NTDs) are a unique feature of Pex1 and Pex6^34^. Other members of the type II AAA-ATPase family, for example Cdc48/p97 or Rix7, contain only a single N-terminal domain, which is nevertheless structurally related to the N-terminal domains of Pex1/Pex6 and known to mediate a plethora of interactions with co-factors and adaptor proteins^35^. The D1 and D2 domains form two stacked heterohexameric rings with a central channel, but in contrast to other type II AAA-ATPases, only the D2 domains of Pex1 and Pex6 are capable to hydrolyze ATP^34,36,37^. Both Pex1 and Pex6 are required for assembly, which occurs in an ATP-dependent manner^38^.

**Figure 1.**
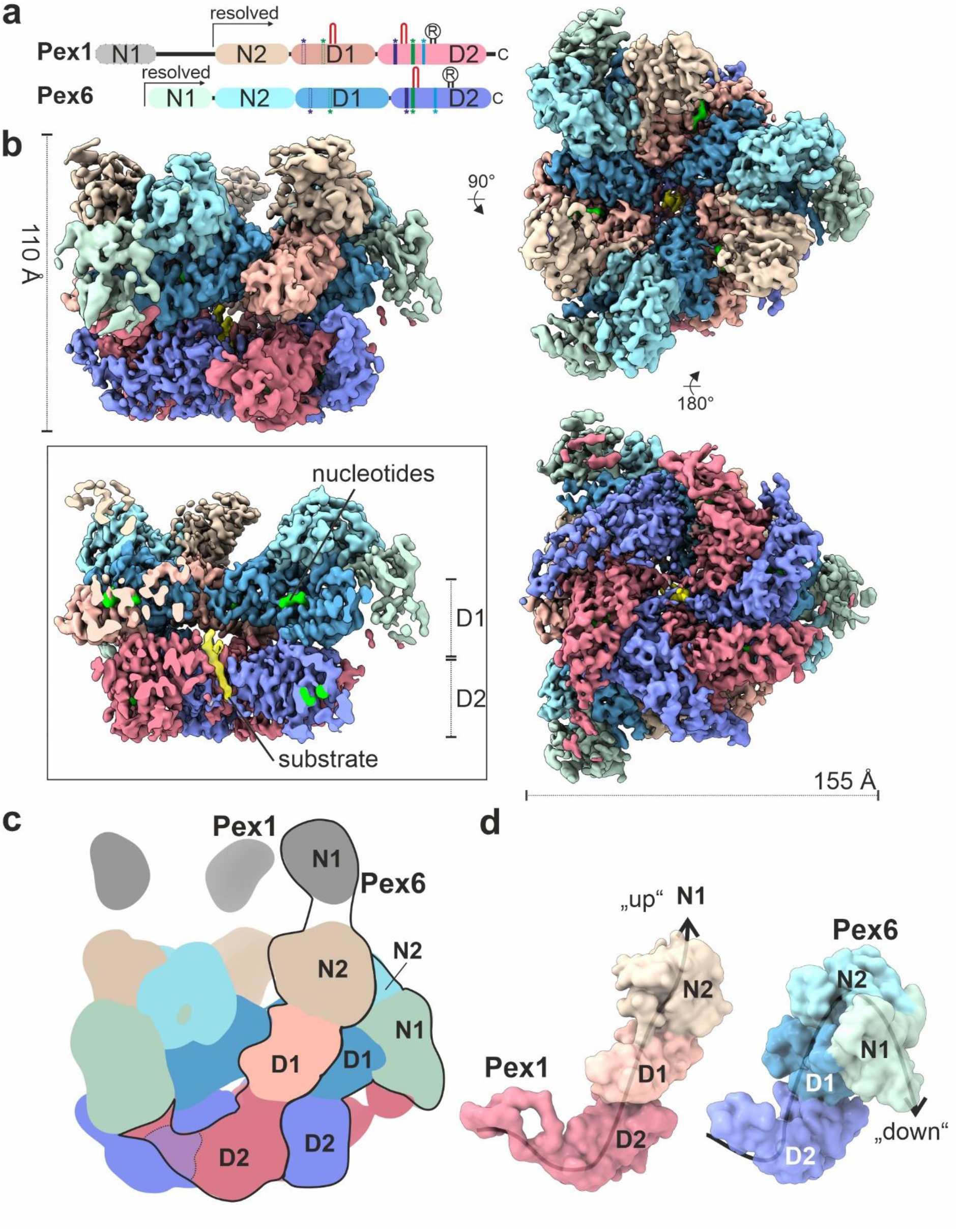
Cryo-EM structure of the peroxisomal ATPase Pex1/Pex6. **a)** Schematic representation of the Pex1 and Pex6 primary structures colored by domain. The color code is maintained throughout the manuscript. Important conserved motifs are indicated: Walker A (dark purple), Walker B (green), inter-subunit signaling (ISS) (cyan), pore loops (red loop) and arginine fingers (R). Dashed boxes (D1) indicate degenerated motifs. **b)** Cryo-EM density map of the better resolved 3D class 3 (Supplementary Figure 2, 3; termed “single-seam” throughout the manuscript) from yeast Pex1-Strep_2_/His-Pex6^(E832Q)^ with each subunit and domain colored as in (a) shown as side, top and bottom view. The inset shows a cut-away view of the cryo-EM structure displaying the central ATPase channel. The density of the substrate and nucleotides are shown in yellow and green, respectively. **c)** Cartoon depicting the overall domain organization of Pex1 and Pex6. The flexible Pex1(N1) domain is not resolved in our cryo-EM density, but shown in an “up” conformation, in accordance with previous low resolution EM studies^37^. **d)** Surfaces of the molecular models of Pex1 and Pex6 shown in the same orientation. Note the characteristic “down”-conformation of Pex6(N1) (right).

In contrast to the D1 and D2 domains, the N-terminal domains of Pex1 and Pex6 do not display high sequence identity to each other and might be responsible for the specific functions of Pex1 and Pex6. For example, the membrane anchor of Pex1/Pex6 (*Sc*Pex15, *Hs*Pex26) binds exclusively to Pex6^21^, whereas the autophagy receptor Atg36 (yeast) binds exclusively to Pex1^32^. The N-terminal regions are not well characterized and might perform additional functions, for example binding to the ubiquitin moiety of Pex5^39^ or association with the ubiquitin hydrolase Ubp15p^40^.

Previous negative stain EM studies in different nucleotide states revealed the general architecture of the heterohexameric Pex1/Pex6 double ring complex, consisting of alternating subunits of Pex1 and Pex6, and suggested large conformational changes in the D1 and D2 rings^23,34,37^. In addition, cryo-EM structures of Pex1/Pex6 in the presence of ADP and ATPγS were determined with an overall resolution between 6.2 Å and 8.8 Å^36^. In contrast to the negative stain EM data, the available cryo-EM structures did not indicate drastic conformational changes and opening of the pore between the ATP*γ*S and ADP states^36^. Despite its essential role in peroxisome biogenesis, import of matrix enzymes and quality control^41^, detailed structural insights into Pex1/Pex6 architecture and mechanism of substrate processing are not yet available.

Numerous high-resolution structures of double-ring ATPases in complex with substrates have now been solved, providing important insights into a rather conserved mechanism of substrate processing through the central pore formed by the D1 and D2 domains^42–48^. However, the coordination between the D1 and D2 ATPase domains in Pex1/Pex6 is complex, involves two different proteins and communication “hubs” between their inactive D1 and active D2 domains. Moreover, it is still unclear whether the underlying molecular mechanisms involve conformational changes of the unique tandem N-terminal domains. For a comprehensive understanding of Pex1/Pex6 function, it is crucial and of considerable biological and pathophysiological interest to visualize Pex1/Pex6 in a “working”, substrate-bound state.

In this study, we captured Pex1/Pex6 from *S. cerevisiae* with an endogenous substrate in the central pore and determined two Pex1/Pex6 cryo-EM structures in different states. These new structures reveal the complex coordinated interplay between Pex1 and Pex6 during substrate processing and allow us to highlight unique features of the molecular motor of peroxisomal receptor recycling.

## Results

In order to obtain high resolution structural information for Pex1/Pex6, we first introduced the E832Q mutation to the Walker B ATPase motif of the D2 domain of Pex6 from *S. cerevisiae* (Pex6_WB). As reported previously, the ATPase activity in the Pex1/Pex6_WB is compromised, despite the presence of WT Pex1^23,34,36,37^. Interestingly, when the same Walker B mutation is introduced to Pex1(D2), the Pex1_WB/Pex6 complex retains some activity^23,34,36,37^. Considering that the D2 domains are responsible for all ATP hydrolysis in Pex1/Pex6^34^, this suggests that the D2 domains of Pex1 and Pex6 control the ATPase activity of the complex in a distinct and possibly highly coordinated manner.

We co-expressed Pex1 WT and Pex6_WB in *S. cerevisiae* and purified the complex by affinity chromatography and size-exclusion chromatography in the presence of a saturating concentration of ATP (Supplementary Fig.1 a,b). We utilized single particle cryo-EM analysis (Supplementary Fig. 1c,d) and finally obtained cryo-EM maps of two distinct states of the complex (Supplementary Fig. 2).

The better resolved conformation (Fig. 1, Class 3 in Supplementary Fig. 2; Supplementary Movie 1) shows Pex1/Pex6_WB in an overall closed conformation with a well-resolved symmetric D1 ring (average resolution 3.7 Å (FSC=0.143); 3.3 Å (FSC=0.5) upon density modification) (Supplementary Fig. 3). The asymmetric D2 ring is resolved to 3.9 Å (3.6 Å (FSC=0.5) upon density modification) (Supplementary Fig. 3). The particle shows a characteristic triangular shape of alternating Pex1 and Pex6 subunits that are arranged around the central ATPase pore. The D1 ring is stacked on top of the D2 ring and crowned by the N-terminal domains (NTDs) (Fig. 1b-d). The D2 ring displays a well-resolved density for five of the ATPase domains and a partially fragmented density for the remaining ATPase domain (4.5 to 7 Å (FSC=0.143), 4 to 5.5 Å (FSC=0.5) upon density modification) (Supplementary Fig. 3). The NTDs of Pex1 and Pex6_WB are well resolved (Fig. 1b), except the N1 domain of Pex1, which is flexibly attached to the rest of the protein via a long linker peptide of 15-20 residues (Fig. 1a,c).

We derived a molecular model of this conformation (Supplementary Fig. 4). Surprisingly, during modeling, we observed a clear density for a peptide of ∼10 residues passing through the central pore of the D2 ring (Fig. 1b; inset). This finding was rather unexpected because we did not add substrate during sample preparation. Similar to other AAA-ATPase structures that have been solved in complex with a substrate^48,48–50^, the Walker B mutation has apparently been beneficial in trapping the substrate, allowing us to capture a snapshot of the translocation process. The density is, however, not sufficiently resolved to allow a clear identification of the polypeptide. The density presumably corresponds to one or a mixture of multiple endogenous substrates that have been trapped in the D2 pore during purification. We were not able to identify substrate density in or above the D1 channel or below the D2 ring. This is consistent with cryo-EM structures of p97 bound to small substrates (Ub_6_), where substrate density was also only resolved in the D2 ring^42^.

### N2 and D1 domains mediate Pex1/Pex6 complex assembly

In this conformation (Fig. 1), the D1 ring is symmetric and all ATPase domains of the D1 ring are bound to ATP (Fig. 2a-c; Supplementary Fig. 5a; Supplementary Fig. 6a-c). Nucleotides bind in a pocket formed at the interface between the large (_L_D1) and small (_S_D1) subdomains of Pex1(D1) and the large subdomain (_L_D1) of Pex6(D1) and vice versa (Fig. 2a).

**Figure 2.**
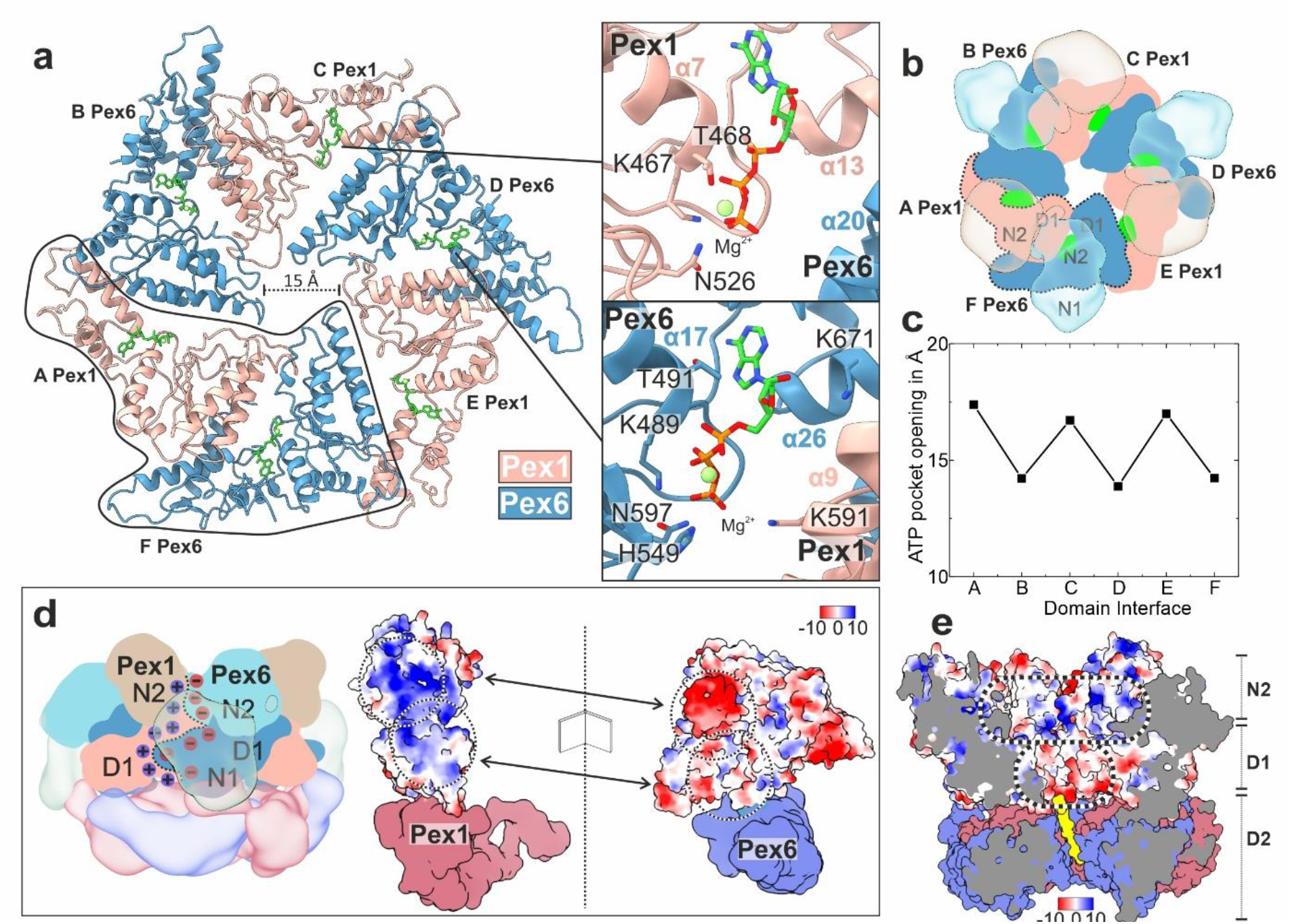
Arrangement of D1 ring and the N-terminal domains. **a)** Top-view of the molecular model of the D1 ring, which resembles a trimer of “tight” Pex6(D1)-Pex1(D1) dimers. Pex1(D1) (beige), Pex6(D1) (cyan), ATP (green). The insets show magnified images of the ATP binding sites at the intra-dimeric Pex6(D1)-Pex1(D1) (upper image) and inter-dimeric Pex1(D1)-Pex6(D1) interface (lower image). **b)** Top-view of the symmetric D1 ring (solid colors) capped by the N-terminal domains (transparent) shown as cartoon. **c)** Plot of the nucleotide binding pocket opening of both interfaces measured as distances between the Cα atom of Pex1 T468 to the clockwise-neighboring Pex6 D582 and Pex6 T491 to Pex1 K563, respectively. **d)** Side view cartoon representation of Pex1/Pex6. The interface between Pex6(N2,D1)/Pex1(N2,D1) dimers (clockwise) is characterized by strong electrostatic interactions. The interface is flipped open to demonstrate the complementary charges at the surface (electrostatic Coulomb potential at pH 7.5). Scalebar kcal/(mol·e) at 298 K. **e)** Surface electrostatic Coulomb potential of the N2-D1 layers at pH 7.5 Positively-and negatively charged surfaces are colored in blue and red, respectively. Scalebar kcal/(mol·e) at 298 K.

The NTDs of Pex1 and Pex6 differ significantly in their conformation relative to the D1 ring (Fig. 1b-d). The N2 domains of Pex1 and Pex6 are closely associated and form a third layer of three Pex6(N2)-Pex1(N2) dimers that crown the D1 ring (Fig. 1c,d, 2b). The flexible Pex1(N1) domain, that is not resolved in our cryo-EM density, is probably located above the N2 layer (Fig. 1c,d). According to previous low resolution EM studies, Pex1(N1) adopts an extreme-”up” conformation (see ^37^). In contrast, Pex6(N1) is well resolved and adopts a characteristic “down” conformation that is coplanar with the D1 ring (Fig. 1c,d). The positions of the NTDs are also similar to the previous EM structures of Pex1/Pex6 determined in the presence of ADP, ATP*γ*S, several nucleotide analogues, and without protein substrate^34,36,37^. This suggests that the overall conformation of the unique tandem NTDs of Pex6 and the N2 domain of Pex1 is independent of nucleotide or substrate binding. In contrast, in p97, the characteristic conformational change of the N-terminal domains from a similar “down” to the “up” conformation is triggered by binding to ATP (D1 domains) and/or binding to substrate and N-terminal cofactors^51–54^. The flexible Pex1(N1) that is not resolved in our cryo-EM structure might play a role in substrate recognition and adopt distinct conformations upon interaction with yet unknown adaptor proteins.

The Pex1(N2) and Pex6(N2) domains are involved in strong interactions to each other and the D1 ring, rendering large independent conformational changes of these domains rather unlikely. In particular, Pex1(N2) and Pex6(N2) form “tight” N2-dimers via strong complementary electrostatic interactions (Fig. 2d). The symmetric D1 ring resembles in general a “trimer” of Pex6(D1)/Pex1(D1) dimers arranged in a clockwise-manner (when viewed from D1 towards D2: top view) (Fig. 2a,b). Each Pex6(D1)/Pex1(D1) dimer is also capped by the “tight” (N2)-dimer, which contributes complementary charges that significantly enforce this interface (Fig. 2d). The intra-dimeric (Pex6(D1)/Pex1(D1) interface is additionally stabilized by ATP (Fig. 2a). Pex6 provides a degenerated Walker-A motif (T491 and K489), a histidine (H549) and a lysine (K671) whereas Pex1 provides an additional lysine (K591) (degenerated Arg-finger) to the heteromeric binding pocket of the nucleotide (Fig. 2a; lower inset). Although the Walker A motif of Pex6 is not archetypal, Pex6(D1) can still bind ATP, which can however not be further hydrolyzed, due to substitutions in the Walker-B and R-finger motifs (Fig. 2a; lower inset).

The Pex1(D1)-Pex6(D1) interface between dimers is less compact and does not involve N-terminal domains (Fig. 2b). This is reflected in the larger opening of the nucleotide binding pocket between Pex1(D1) and Pex6(D1) (Fig. 2a, upper inset), when compared to the respective pocket at the intra-dimeric interface (Fig. 2a, lower inset). In this case, ATP interacts with residues T468 and K467 (Walker A-like), and N526 (Walker B-like) that are contributed by Pex1(D1). Surprisingly, Pex6(D1) does not interact with the ATP bound to Pex1, due to the large distance of the involved domains and the deletion of the R-finger motif (Fig. 1a, 2a). Our data thus suggest that ATP does not directly contribute to the inter-dimeric D1 contact but stabilizes instead the small ATPase domain Pex1(_S_D1) that is involved in the respective interactions.

Despite high structural similarity, Pex1(_S_D1) and Pex6(_S_D1) differ significantly in size (Supplementary Fig. 7a,b). Pex6(_S_D1) is 20 residues longer than Pex1(_S_D1) (Supplementary Fig.7b; Pex6 helix α26 and α27). This feature dictates the compactness of the interface between _S_D1 and _L_D1. The loose interface between dimers (Pex1(_S_D1)→Pex6(_L_D1)) involves the “shorter” Pex1(_S_D1), whereas the compact dimeric interface (Pex6(_S_D1)→Pex1(_L_D1)) involves the “longer” Pex6(_S_D1) (Fig. 2a; Supplementary Fig. 7a). In Pex6(_S_D1)→Pex1(_L_D1) (compact dimeric interface; (Fig. 2a; Supplementary Fig. 7a), M679 of the prolonged helix α26 of Pex6(_S_D1) docks into a hydrophobic pocket presented by Pex1(_L_D1) (Supplementary Fig. 7c). Furthermore, the prolonged helix α26 of Pex6(_S_D1) displays a negatively charged pocket that allows binding of the conserved H592 of Pex1(_L_D1) (Supplementary Fig. 7c). In contrast, at the less compact Pex1(_S_D1)→Pex6(_L_D1) interface between dimers, the analogous H609 of Pex6(_L_D1) is not involved in any interactions (Fig. 2a; Supplementary Fig. 7d, yellow highlight), due to the deletion in the opposite helix α14 of Pex1(_S_D1). This allows the analogous hydrophobic pocket of Pex1(_L_D1) to click further downstream into the conserved Y654 of the shorter helix α13 of Pex1(_S_D1) (Supplementary Fig. 7d). The consequence is a larger angle between Pex1(_S_D1) and Pex6(_L_D1) and thus a less compact interface between two adjacent dimers. In summary, the deletion in helix α13 and α14 of Pex1(_S_D1) determines the “trimer-of-dimers” arrangement in the D1 ring of Pex1/Pex6 complex.

Each D1 heterodimer is further capped by the associated NTDs, which further results in the characteristic triangular shape of Pex1/Pex6 when viewed from the top (Fig. 1b, 2b). The resulting central pore of the D1 ring has a diameter of approximately 15 Å (Fig. 2a; Supplementary Fig. 6d). The D1 channel, however, lacks substrate density (Fig. 1b). This renders tight interactions between D1 and the substrate unlikely. Interestingly, the “crown” assembled by three N2 heterodimers forms a positively charged funnel leading to the negatively charged “mouth” of the D1 ring (Fig. 2e).

### The Pex1/Pex6 D2 ring motor processing substrate

Instead of a planar pseudo-symmetric arrangement, as observed in the previous reconstructions of Pex1/Pex6 in the presence of ADP and ATPγS^36^, the AAA-ATPase domains of the D2 ring of substrate bound Pex1/Pex6 assemble into an asymmetric right-handed spiral staircase (Fig. 3), similar to other ATPases^55,56^. The substrate polypeptide chain presumably stabilized this conformation. Similar to the D1 ring, Pex1(D2) and Pex6(D2) are also organized in pairs (Fig. 3a). The Pex6(_S_D2)→Pex1(_L_D2) dimeric interface is indeed stronger than the Pex1(_S_D2)→ Pex6(_L_D2) interface between the D2 dimers (Supplementary Fig. 8).

**Figure 3.**
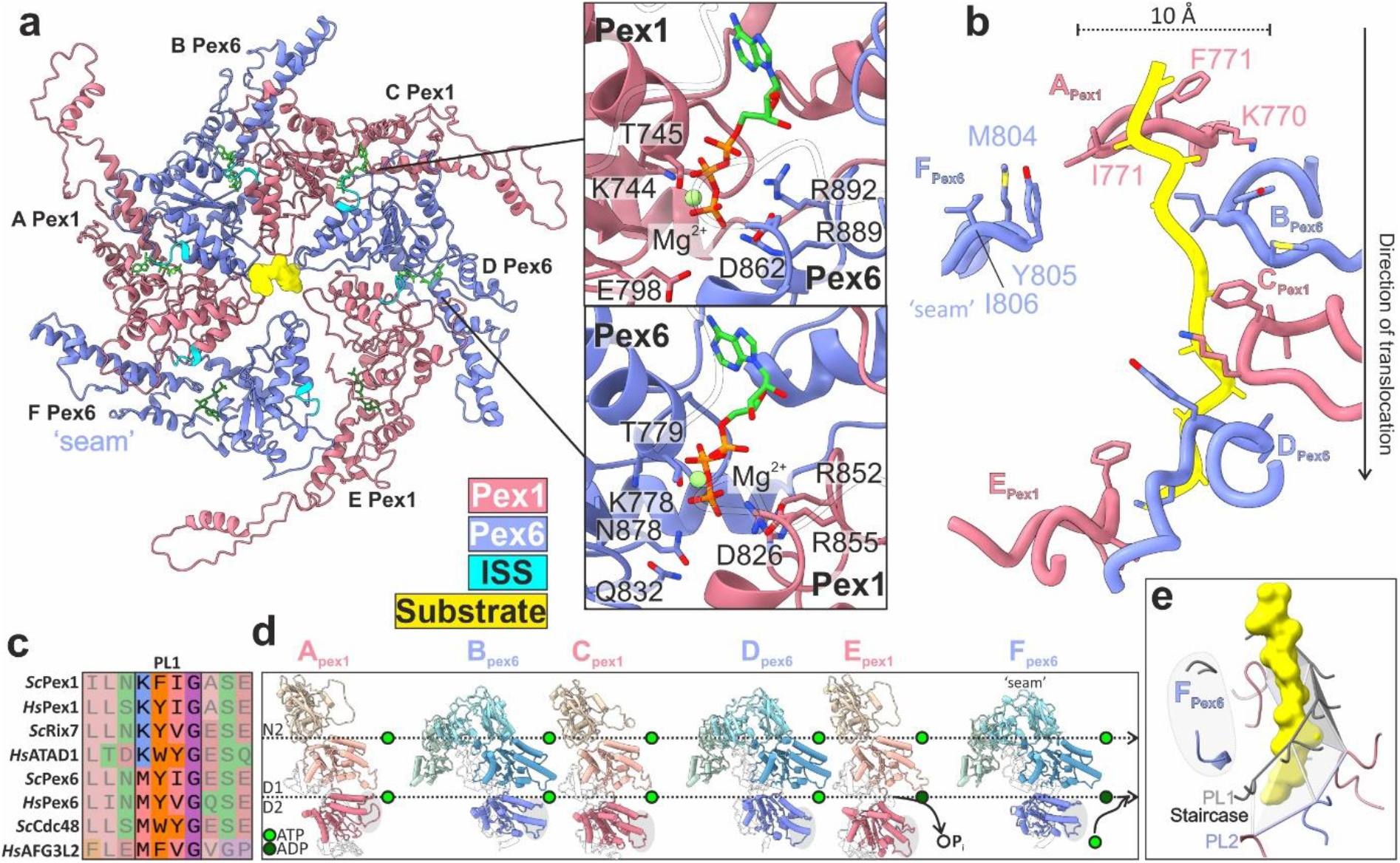
Substrate engagement in the active Pex1/Pex6 D2 channel. **a)** Top view of the substrate-bound Pex1/Pex6 D2 ring. Pex1(D1) (coral), Pex6(D2) purple, ISS motif (cyan), nucleotides (green) and substrate (yellow). The insets show magnified images of the ATP binding sites at the Pex1(D2)-Pex6(D2) (upper image) and Pe6 (D2)-Pex1(D2) interface (lower image) of substrate engaged subunits C and D. **b)** Residues of pore loops 1 contacting the substrate. **c)** Multiple alignments of pore loop 1 sequences of D2 domains from several AAA-ATPases including Pex1, Pex6, Rix7, p97/Cdc48, AFG3L2 and ATAD1. **d)** Pex1 and Pex6 subunits shown side-by-side, aligned along their D1 domain. Note the sequential tilt of the D2 domain relative to the D1, which is highlighted by a dashed line above each domain. The catalytically “dead” D1 domains remain ATP-bound. **e)** Spiral arrangement of pore loops 2. Pore loops 1 are shown in grey and substrate in yellow. The pore loops 2 of subunits Pex1(A) to Pex6(E) (PL2) form a second spiral staircase running below and parallel to the spiral staircase formed by the pore loops 1 (PL1). The pore loops of the “seam” subunit Pex6(F) (highlighted in gray) are displaced significantly from the spiral staircase.

The D1 and D2 rings are rather flexibly connected via the D1→D2 linker peptides and the protrusion domains of Pex1(D2)(helix α28) and Pex6(D2)(helix α38) (Supplementary Fig. 9a-b). The protrusion domain of Pex6(D2) interacts with Pex1(D1) within a subunit dimer (Supplementary Fig. 9c, lower panel), whereas the more flexible protrusion domain of Pex1(D2) (Supplementary Fig. 9b,c) interacts with the neighboring Pex6(N1), linking thereby two adjacent subunit dimers (Supplementary Fig. 9c, upper panel).

Within the D2 ring, the interfaces between the D2 subdomains (Pex1(D2)/Pex6(D2) (between dimers) and Pex6(D2)/Pex1(D2) (intra-dimer)) are further stabilized by nucleotides. The binding pockets are formed by conserved Walker A and Walker B residues interacting with a pair of arginine-finger residues of the clockwise neighboring subunit (Pex1: R852, R855; Pex6:R889, R892) (Figure 3a; Supplementary Fig. 5a, lower panels). Mutation of the Arg-finger residues Pex1(R852) and Pex6(R889) to lysine results in 100% and 80% inhibitions of the ATPase activity of the complex, respectively^34,37^.

The substrate is threaded into a right-handed spiral staircase formed by pore loops 1 of Pex1(D2) and Pex6(D2). The central pore of the spiral staircase has a diameter of ∼10 Å (Fig. 3a,b). We were able to model a continuous poly-alanine polypeptide backbone of 10 amino acids into the respective density. The translocating peptide was modelled with N→C directionality (Fig. 3b). This is consistent with the translocation direction of other substrate-engaged cryo-EM structures of ATPases, exhibiting the same characteristic right-handed spiral staircase arrangement of pore loops 1, as observed in our structure^42–45,47,57^. The pore loops 1 of chain A (Pex1) and E (Pex1) occupy the highest and the lowest position of the spiral (Fig. 3b, grey highlight in 3d). D2 of chain F (Pex6) does not directly contact the substrate and does not follow the spiral arrangement (“seam” subunit) (Fig. 3a,b). We therefore define this conformation of the heterohexamer as the “single-seam” state. The substrate engaged pore loops 1 of D2 domains Pex1(A,C,E) and Pex6(B,D) are related to each other by a ∼60° rotation (axis of Pex1-helix α_22_ to Pex6-helix α_30_) and a distance of 8-9 Å (Pex1 residue 771 Cα to Pex6 residue 805 Cα). The D2 pore loops 1 of Pex1 and Pex6 contain only a single aromatic residue in a K**F**I-or M**Y**I-motif, respectively. The aromatic residue (Pex1 F771; Pex6 Y805) and the isoleucine of the involved pore loops 1 bind to the substrate every two amino acids in a “forceps”-like manner (Fig. 3b). The preceding methionine (M804) of Pex6(D2) and lysine (K770) of Pex1(D2) mediate the communication between the adjacent pore loops and stabilize their arrangement (Fig. 3b). M804 of Pex6 contacts F771 of the adjacent clockwise Pex1(D2), whereas K770 of Pex1 forms π-cation interactions with the aromatic Y805 of the adjacent clockwise Pex6(D2) (Fig. 3b). The residue prior to the aromatic in pore loop 1 is in general suggested to have a significant role on substrate translocation^48^. The Pex1(D2) pore loop 1, with lysine forming π-cation interactions, is similar to the staircase of Rix7 and ATAD1 (Fig. 3c). The M804 of the Pex6(D2) pore loop 1 is rather engaged in van-der-Waals interactions, similar for example to Cdc48 or AFG3L2 (Fig. 3c). The D2 ring thus features two types of pore loops 1 with distinct characteristics. Whether and how these hybrid features of the spiral-staircase have an effect on fine-tuning substrate translocation and Pex1-Pex6 coordination during this process requires further investigation.

The four D2 domains at the top of the spiral are ATP-bound (Pex1(A), Pex6(B), Pex1(C), Pex6(D) (Figure 3b,d; Supplementary Figure 5a; lower insets). Assessing the nucleotide in the binding pocket of the “lowest” Pex1(E) and “seam” subunit Pex6(F) (Fig. 3b,d) based on the cryo-EM density alone, was not possible (Supplementary Fig. 5a). We therefore measured the buried surface area between adjacent protomers and distances between Walker A and arginine-fingers of the associated protomers (Supplementary Fig. 5c-e). The extent of nucleotide binding pocket opening is a reliable and well-established indicator of the nucleotide status of the respective subunit^44,48,58^. This analysis revealed that both the “bottom” Pex1(E) subunit and the disengaged “seam” Pex6(F) D2 domains are in an ADP-bound state, as previously described for other AAA-ATPases (Fig. 3c,e, Supplementary Fig. 5a,c-e)^42,45,59^. The position of the pore loops 1 in the spiral staircase thus highly depends on the nucleotide state of the respective subunit (Fig. 3b,d).

The pore loops 2 of engaged Pex1(D2) and Pex6(D2), except pore loop 2 of “seam” Pex6(F)), surround the substrate and form a second spiral staircase, below the spiral staircase formed by the pore loops 1 (Fig. 3e). The pore loops 2 are less well resolved, lack aromatic residues and are not tightly associated with the substrate. Similar to other AAA-ATPases^55^, they provide polar and negatively charged residues facing the substrate. This might support the translocation process by stabilizing the backbone of the substrate and preventing its refolding during threading.

### Pex1 and Pex6 are highly coordinated

The second cryo-EM structure of substrate-bound Pex1/Pex6_WB (Supplementary Fig. 10) shows an intriguing open state where both the D1 and D2 rings of Pex1/Pex6 have an asymmetric configuration (Fig. 4a). One of the Pex6(D1)/Pex1(D1) dimers thereby swings out. This results in an open central D1 channel with a diameter of 20 Å, approximately 5 Å larger than the symmetric D1 ring in the “single-seam” state (Fig. 4a; Supplementary Fig. 6a,d-f). Interestingly, the outward Pex6(D1)/Pex1(D1) dimer is stacked on top of a compact characteristic pair of dislocated seam domains of the D2 ring Pex6(D2)(E)/Pex1(D2)(F) (Figure 4a,b). We therefore term this conformation as the “twin-seam” state of the complex.

**Figure 4.**
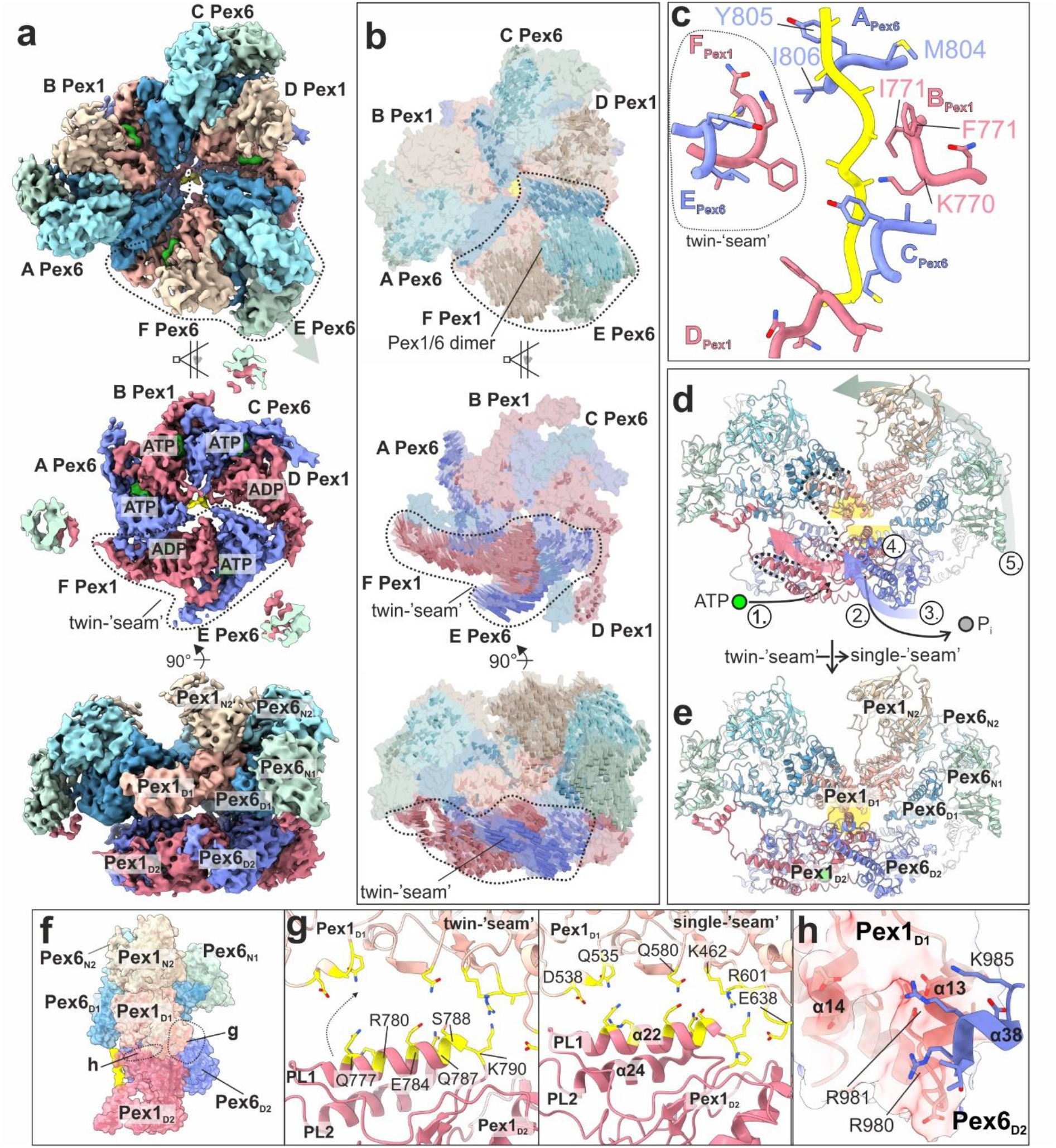
Architecture of Pex1/Pex6 “twin-seam” state. **a)** Cryo-EM structure of “twin-seam” state of yeast Pex1/Pex6_WB shown as top (upper panel), cut-(middle panel) and side view (lower panel). Individual domains and the substrate are colored as in Figure 1(a). Note that the “twin-seam” subunits (dashed line) are dislocated from the substrate. **b)** Molecular surface of Pex1/Pex6 with vectors indicating the putative direction of motion during the conformational switch from the “twin-seam” to the “single-seam” state. **c)** The pore loop 1 staircase with the twin-seam subunit being disengaged from the substrate. Residues of an individual loop of Pex1 and Pex6 are labeled. **d-e)** Side view of the molecular model of Pex1/Pex6 dimer in the “twin-seam” (d) and “single-seam” (e) state. Possible sequential steps of the conformational switch are indicated: 1) ATP-binding between dimers 2) ATP-hydrolysis within the “twin-seam” dimer 3) rotation 4) D2→D1 docking, and 5) D1, N swinging. The “communication hub” between the D2 and D1 ring is highlighted in yellow. **f)** Side view of the Pex1/Pex6 dimer with interaction hubs being highlighted (Pex1(D2)/Pex1(D1)(g); Pex6(D2)/Pex1(D1)(h)). **g)** Pex1(D2)/Pex1(D1) interface in “twin-seam” (left panel) and “single-seam” state (right panel). This interface involves the Pex1(D2) helix α22 located downstream of pore loop 1. During the twin-seam (left panel) to the single-seam (right panel) transition, the helix moves, establishing new interactions. Residues are shown as sticks. **h)** Pex6(D2)/Pex1(D1) interface (single-seam state). Surface of the negatively charged groove of Pex1(D1) colored according to the electrostatic potential. Interacting residues of Pex6(D2) are shown as sticks.

Note that all Pex1(D1) and Pex6(D1) domains of the D1 ring remain bound to ATP (Supplementary Fig. 5b; upper panels), but the deletion of the R-finger motif in Pex6(D1) apparently renders the inter-dimeric interfaces flexible. This allows even greater distances between the outward dimer and adjacent subunits (Supplementary Fig. 6a-c). Substrate density is again limited to the D2 channel. In the “twin-seam” state, only four alternating Pex6(D2) and Pex1(D2) domains (Pex6(A), Pex1(B), Pex6(C), Pex1(D)) form the substrate-interacting spiral staircase of pore loops 1 (Fig. 4c).

The “twin-seam” dimer Pex6(E)/Pex1(F) is dislocated from the central channel (Fig.4a). The pore loops 1 of the “seam” subunits are displaced significantly from the spiral staircase (Fig. 4c). This characteristic “twin-seam” pair is positioned between the lowest (Pex1(D)) and the highest position of the staircase (Pex6(A)) (Fig. 4c).

The loose interfaces of the “twin-seam” dimer to the neighboring D2 domains are profoundly different from the close contacts between the other D2 domains, including a rather “tight” interface between the two D2 domains within the “twin-seam” dimer (Fig. 4a; Supplementary Fig. 5c-e). The nucleotide binding site shared by the Pex1 (D2) “seam” (F) and the clockwise (from top) Pex6(D2) (A) is not well defined, thus it is either empty or ADP-bound (Supplementary Fig. 5b; lower panels (F); Supplementary Fig. 5c-e). Subsequent subunits contacting the substrate (Supplementary Fig. 5b; lower panels, (A,B,C), Supplementary Fig. 5c-e) exhibit strong density in the nucleotide binding pocket and tight interfaces between the respective motifs, suggesting that they are bound to ATP. The nucleotide binding site between the Pex6(D2) “seam” and the counter-clockwise Pex1(D2) (Supplementary Fig. 5b lower panels, (D); Supplementary Fig. 5c), show density suggestive of the presence of a nucleotide. The wide spacing between the involved motifs suggests that ADP is bound to this pocket (Fig. 4a; Supplementary Fig. 5c-e). The lowermost D2 domain of the staircase is thus bound to ADP and the “twin-seam” dimer is released from the staircase (Fig. 4c). Intriguingly, our analysis suggests that the nucleotide binding pocket (E) between the two “seam” domains remains bound to ATP (Supplementary Fig. 5b-e).

Taking into account that ATP-hydrolysis occurs in an anti-clockwise manner (when viewed from top), we compared the “twin”-(Fig. 4a) and the “single”-seam (Fig. 1b) structures and visualized the direction of motion between both states (Fig. 4b). It is interesting to note that both Pex1(D2) and Pex6(D2) of the “twin-seam” dimer move synchronously as a unit towards the preceding Pex6(D2) indicated as arrows (Fig. 4b). We assume that this drastic conformational change towards the “single-seam” conformation is triggered upon binding of ATP to the binding pocket (F) of the “twin-seam” (Supplementary Fig. 5c; lower panel). Parallel to the clockwise rotation (viewed from top) of the “twin-seam,” (Fig. 4b, middle panel) the respective Pex6(N2/D1)/Pex1(N2/D1)-dimer swings in toward the central channel (Fig. 4b, upper panel), resulting in the symmetric D1 ring of the “single seam” structure (Fig. 1b). Thus, there is a coordinated communication between the two rings, although the D1 ring is catalytically dead with respect to ATP hydrolysis.

Upon careful inspection of the conformational changes between the two states, we identified a major “communication-hub” between the D2 domain of Pex6 and the D1 domain of Pex1, within a Pex1/Pex6 dimer (Fig. 4d-e). Binding of ATP to the binding pocket of (F) triggers the rotation of the “twin-seam”, so that “seam” Pex1(D2)(F) engages the substrate at the highest position of the spiral (Fig. 4a-e). This results in binding of pore loop 1 of Pex1(D2) (F) to the substrate, upwards movement of Pex1-helix_α22_ (downstream helix of pore loop 1), which in turn establishes new strong interactions, pulling on Pex1(_L_D1) (Fig. 4d-g, Supplementary Movie 2,3). In parallel, the rotation of the Pex6(D2)/Pex1(D2) “twin-seam” dimer (Fig. 4b, middle panel, Supplementary Movie 2), pushes the Pex1(_S_D1) upward. This is mediated via the Pex6 protrusion helix α38 (Fig. 4d,f,h; Supplementary Fig. 9c(ii)). In particular, three positively charged residues of Pex6(D2)-helix_α38_ (R980, R981, K985) dock into this groove and push Pex1(_S_D1) upward (Fig. 4h, Supplementary Movie 2; Supplementary Fig. 9c(ii)).

These conformational changes are propagated to the Pex6(D1)/Pex1(/D1) dimer on top of the “twin-seam”, that swings toward the central channel and pulls the associated Pex6(N2) domain along that direction (Fig. 4b,d,e). However, it should be noted that the Pex6(N2) thereby retains its “down” orientation and remains connected to the Pex1(D2) protrusion domain. In summary, the described Pex6(D2)/Pex1(D1) pivot (Fig. 4d-e, yellow highlight) translates the rotation of the D2 “twin-seam” during the ATP hydrolysis cycle (Fig. 4b, middle panel) into a coordinated large inward tilting of the associated D1 dimer and N-terminal domains (Fig. 4b, upper panel).

## Discussion

Cryo-EM structures of numerous unfoldases revealed a highly conserved hand-over-hand mechanism of processive threading of diverse substrates along the central channel of AAA+ family members^42–50,56,57^. Pex1/Pex6 is a less well characterized type II AAA-ATPase that plays a crucial role in recycling of peroxisomal receptors. About 65% of PBDs (including Zellweger) are related to mutations of the human Pex1/Pex6 complex^4,5,32,33,60^.

Pex1/Pex6 is the only known double-ring heterohexamer of the type II AAA+ family, with both subunits capped by a unique pair of N-terminal domains (Fig. 1). The D1 ring of the heterohexamer is catalytically inactive^34,37,61^ and ATP hydrolysis is limited in the D2 ring. Pex1 and Pex6 are arranged in a parallel manner to form oblique dimers that are capped by a tight Pex1(N2)/Pex6(N2) dimer, stabilized by strong electrostatic interactions (Fig. 2d). Our structures suggest the Pex1/Pex6 dimer is thus the building block of the heterohexameric assembly, throughout all layers of the complex, including the D2 ring. ATP plays a crucial role in assembly of the hexamer. All D1 binding pockets are occupied (Fig. 2) and previous studies have shown that the complex does not assemble in absence of ATP^62^.

The molecular mechanism of how Pex1 and Pex6 work together to pull the peroxisomal receptor out of the peroxisomal membrane for another round of import has remained unknown. Our data substantially extend our knowledge about the molecular architecture and function of the yeast Pex1/Pex6 complex.

We used a Pex6 Walker B mutant (E832Q) and “trapped” the heterohexamer processing an endogenous substrate along the central channel formed by the active D2 ring. Our cryo-EM structures establish that the Pex1/Pex6 heterohexamer is at first glance a “canonical” unfoldase engaging the substrate via a spiral staircase of conserved pore loops 1 along the central pore of the D2 ring^42–45,48,56^. Similar to other ATPases, Pex1/Pex6 operates in processive manner, with conserved pore loops 1 engaging the substrate and pulling it towards the exit of the D2 channel^42–45,48,56^. The N-terminal domains Pex1(N2), Pex6(N1), and Pex6(N2) do not rearrange during the ATP hydrolysis cycle, as is the case with p97^51,52,63^. The flexible Pex1(N1) domain is not resolved in our cryo-EM structures and might be involved in interactions with the substrate and/or yet unknown co-factor or adaptor proteins^64,65^.

We were able to localize the substrate in the D2 channel but due to limited resolution, we were not able to reveal its identity. The Pex1/Pex6 complex has been previously proposed to solely target ubiquitinated Pex5, but more recently it has been shown that the complex is also capable to process its membrane anchor Pex15^23^, which binds to the N2 domain of Pex6 ^23,66^ or the pexophagy inducing receptor Atg36^32^. We were not able to identify the substrate density in the D1 pore, suggesting that the substrate and the D1 domains might not establish stable contacts. Interestingly, for Cdc48/p97, polyubiquitinated Ub-Eos (Ub_n_-Eos) was observed both in the D1 and D2 rings^42,45^. In contrast, the smaller substrate hexa-ubiquitin (Ub6) was resolved only in the D2 ring^42^, similarly to our cryo-EM structures.

The fact that the hexamer is built of alternating Pex1/Pex6 heterodimers has a major impact on the mechanism of substrate processing. Our study revealed two distinct functional states of substrate engaged Pex1/Pex6. In the better resolved state (dubbed “single seam”), the inactive D1 ring is symmetric (Fig. 2) and resembles a trimer of Pex6(D1)/Pex1(D1) dimers. The catalytically active D2 shows a spiral arrangement that drives translocation (Fig. 3b,e), as previously described for other unfoldases^42– 45,48,56^. Aromatic residues of five alternating Pex6(D2)-and Pex1(D2)-pore loops 1 arranged in a spiral staircase engage the substrate (every two amino acids of the substrate) (Fig. 3b). The sixth loop is dislocated and thus, does not contact the substrate. This loop is positioned between the lowest and the highest position of the spiral staircase (“seam”). Interestingly, Pex1(D2) and Pex6(D2) provide distinct pore loops 1 with different properties regarding substrate-grip and inter-pore-loop 1 coordination (Fig. 3b).

The second “twin-seam” state reveals two “seam” D2 domains forming a characteristic Pex6(D2)/Pex1(D2) pair, that is detached from the spiral (Fig. 4c). The associated Pex6(D1)/Pex1(D1) dimer is tilted outward and the D1 channel is widened (Fig. 4a; Supplementary Fig. 6e,f). Intriguingly, the pore loops 1 of the “twin-seam” in the D2 ring are close together and almost at identical heights, which, at least to our knowledge, has not been observed for canonical ATPases bound to substrate (Fig. 4c). Our data clearly suggest that Pex1(D2) and Pex6(D2) function in concert with each other.

We propose a processive threading and inter-ring communication mechanism, shown schematically in Fig. 5, which involves Pex1(D2)/Pex6(D2) pairs and allosteric force propagation to the D1 ring. In the first step of the cycle, two Pex6(D2)/Pex1(D2) dimers are bound to the substrate (Figure 5b, step i). The characteristic D2 “twin-seam” dimer is dislocated from the central channel of the complex. The associated Pex6(D1)/Pex1(D1) dimer on top of the D2 “seam-dimer” is aligned to this arrangement and is also dislocated from the central D1 channel (Fig. 5a, step i). This results in an “open” conformation of the complex (Supplementary Fig. 6e).

To explain the transition from the “twin seam” to the “single seam” state, we propose that once the Pex1(D2) “seam” (F in Fig. 5b,c, step i) binds ATP and engages the substrate at the top of the spiral (moves to position A in Fig. 5b,c, step ii), the second domain of the D2 “seam” dimer, Pex6(D2)(E), follows towards the highest position of the staircase. However, it still does not contact the substrate and remains as a “single seam” subunit (becomes F in Fig. 5b,c, step ii).This is the step where the pair separates transiently during the ATP hydrolysis cycle. However, due to the “clicking” of the first pore 1 loop of the Pex1(D2)(A)/Pex1(D2)(F) dimer on the spiral, there are steric interactions at a “communication” hub between Pex6(D2) and Pex1(D1) within the dimer (Fig. 4d). This triggers a swing-in of the Pex1(D1)/Pex6(D1) dimer stacked on the D2 “twin seam” and associated N-terminal domains toward the central channel, resulting in a “closed” conformation of the D1 ring (Fig. 5a, step ii, Supplementary Fig. 6d). Pex1(D2)(E) (Fig. 5b,c, step ii) subsequently arrives at the bottom of the spiral. However, in the next step, this domain is not released from the spiral and does not become a “single seam” protomer, as would be the case in a canonical sequential ATPase cycle when hydrolysis at subunit D leads to the release of subunit E.

**Figure 5.**
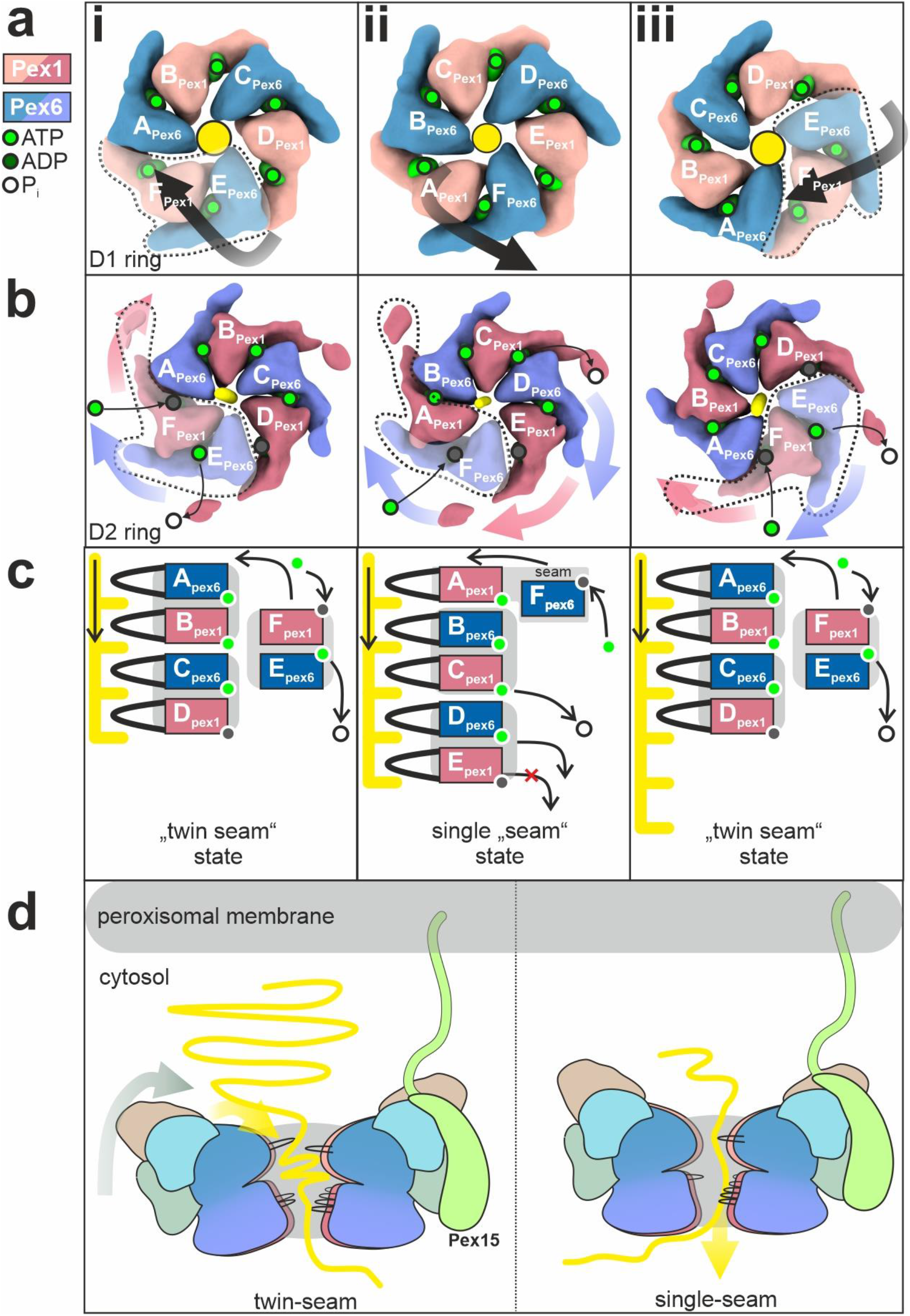
Proposed model of Pex1/Pex6 assembly and substrate translocation. **a-c)** Schematic of proposed Pex1/Pex6 model for ATP-fueled substrate processing. Top view of the D1 (a) and D2 ring (b) and schema of the pore1-loop arrangement (c) in “twin-seam”(i)→”single-seam”(ii)→”twin-seam”(iii) conformational transitions. The substrate is shown in yellow. The subunit occupying the highest position of the right-handed spiral staircase at the respective state, is defined as subunit “A”. Pex1(D2) and Pex6(D2) are alternating in the D2 ring, thus subunit A can be either Pex1 or Pex6. Note that the ATPase does not rotate during the ATPase cycle, rather the ATP is hydrolyzed in a counterclockwise manner. Also note that both Pex6 and Pex1 form dimers both in D1 (a) and D2 (b) rings. Pex1(D2)/Pex6(D2) heterodimers are highly coordinated in processing the substrate. The D1 ring is inactive, but the Pex1(D1)/Pex6(D1) heterodimer tilts inwards in the “twin-seam”(i)→”single-seam”(ii) transition, whereas it swings out in “single-seam”(ii)→”twin-seam”(iii) transition. **d)** The Pex1/Pex6 heterohexamer binds to Pex15 at the peroxisomal membrane, with the “mouth” of the D1 ring facing the peroxisome. In the “twin-seam” state, the D1 dimer opposing the spiral staircase (stacked on top of the twin seam), is tilted outward in an open conformation. Triggered by the power stroke in the D2 ring, the D1 dimer tilts inward and “pushes” the substrate towards the opening of the spiral staircase, supporting thereby substrate unfolding.

Instead, it remains tightly bound to its dimer partner Pex6(D2)(D) (Fig. 5b,c, step ii) because the nucleotide at position D (Pex6(D2)) is not hydrolyzed. Translocation of the substrate comes to a halt because the spiral cannot move (Fig. 5b,c, step ii). The ATPase skips this position and instead hydrolyzes the nucleotide at position C (Pex6(D2)), which is located between the dimers (Fig. 5b,c, step ii). The ATP hydrolysis at this position, provides thus a power stroke to release the Pex6(D2)(D)/ Pex1(D2)(E) dimer at the bottom of the spiral, which simultaneously allows the engagement of Pex6(D2)(F) (formerly “single seam”) at the top of the spiral after it has bound to ATP (Fig.5b,c, step ii). The release of the “twin-seam” is propagated to the D1 ring and the associated D1 dimer swings out (Fig. 5a, step iii). Once the “twin seam” is released, hydrolysis is now possible at position E due to the loosening of the dimer interface (Fig. 5b,c, step iii). This in turn triggers the “twin seam”→”single seam” transition. In summary, our data cannot be reconciled with the traditional “hand-over-hand” model. The ATPase cycle of Pex1/Pex6 involves the release of “two hands” as the uncoupling of Pex6(D2) from the substrate is highly coordinated with Pex1(D2).

To understand the underlying coordination mechanism of “twin-seam” release, we examined the conformation of the intersubunit signaling motif (ISS) in the “single-seam” state (Fig. 5c, step i). Recent studies on substrate-bound p97 revealed a conformation of ISS capable of controlling the nucleotide status of the adjacent subunit by projecting into the nucleotide binding pocket of the adjacent subunit and blocking ATP hydrolysis^42^. ATP hydrolysis occurs only once the ISS motif is retracted (Supplementary Fig. 11a)^42^. Thus, in p97, the ADP state of subunit E at the bottom of the spiral is communicated to the immediately adjacent subunit, triggering ISS-retraction and allowing the staircase to progress sequentially (Supplementary Fig. 11a)^42^. In Pex1/Pex6 “single-seam” state (Supplementary Fig. 11b), however, the D2 domains are organized as tight dimers and this signal is not passed on to the directly neighboring subunit, but two subunits further. On the one hand, the ISS motif of Pex1(E) remains inserted and prevents the hydrolysis of the directly neighboring Pex6(D). On the other hand, the ISS motif of Pex6(D) is in an intermediate conformation and is almost retracted from the nucleotide site of Pex1(C), which is located at the dimeric interface. Therefore, as expected, this site is predisposed to ATP hydrolysis, which explains the release of the “twin seam” Pex6(D)/Pex1(E) (Fig.5c, step ii) instead of Pex1(E) as a “single seam.” Most importantly, it is only after the release of the dimer (Fig. 5c, step iii) that the dimeric interface is loosened and the ISS motif of Pex1(F) is retracted, allowing hydrolysis at the dimeric interface (Supplementary Fig. 11c,d-e) (Pex6(E)) and continuation of the ATPase cycle. In summary, our data suggest that the ISS-motif is critical for the coordination of Pex1/Pex6 pairs and release of the “twin-seam”.

However, an important question remains unanswered: “What is the functional significance of the opening of the D1 ring, powered by the D2 ATPase cycle?” A possible answer is that the “swinging” might be helpful to position and “push” the substrate towards the “mouth” of the central channel, thereby supporting the unfolding of the substrate (Fig. 5c). It should be emphasized that Pex1/Pex6 is so far lacking co-factors, known to facilitate correct orientation for substrate binding in other AAA-ATPases. Furthermore, the swing occurs in the D1 dimer, which opposes the spiral staircase in the underlying D2 ring (Supplementary Fig. 6f). In the absence of cofactors, inward movement and pore closure might thus be required to position the incoming substrate at the opening of the pore (Fig. 5d), exerting forces that support unfolding and making the ATPase more efficient.

The role of Pex1(N1) in this process remains unclear. This domain is flexible and not resolved in the cryo-EM density. It is expected to bind substrates or co-factors (similar to the N-domains of p97), but it is yet unclear if it binds ubiquitin^64^. It is also possible that the D1 tilting propagates forces to interaction partners. For example, it might also affect the membrane anchor of Pex15 via Pex6(N2) (Fig. 5d), which links the peroxisomal exportomer with the importomer and *vice versa*^14,27,67^.

Recent cryo-EM data suggest that the peroxisomal Ub ligase complex, required for peroxisomal receptor recycling, might form a retro translocation channel at the peroxisomal membrane^19^. We expect that Pex1/Pex6 dimers assemble to hexamers near the peroxisomal membrane and bind via the Pex6(N2) domain to the elongated membrane anchor Pex15 (Fig. 5c)^23,41^. Pex15 would then orient the D1 ring facing towards the peroxisomal membrane, possibly in close proximity to the ligase complex. The ubiquitinated receptor Pex5 would then be taken up by the Pex1/Pex6 complex and pulled back into the cytosol^18^ (Fig. 5c, yellow line). However, it is still unclear how the ubiquitinated Pex5 receptor is delivered to the ATPase channel and whether other cofactors and/or Pex1(N1) are involved.

In summary, this study reveals the architecture of the molecular motor of peroxisomal receptor recycling and suggests a mechanism of substrate translocation and utilizing communication between D1 and D2 rings, with fundamental differences to other type II AAA-ATPases. Our data establish a solid foundation for future studies towards understanding the complex interplay of the peroxisomal exportomer and importomer machinery.

## Acknowledgements

This project has been supported by FOR 1905 (PerTrans consortium) GA 2519/1-2 to

C.G. and ER 178/7-2 to R.E. from the German Research Foundation (DFG). C.G. acknowledges support from the research consortium CRC1430 (DFG). The cryo-EM dataset was processed at the Palma II HPC (INST 211/667-1) of the University of Münster. We acknowledge technical support by the HPC team of the WWU IT, especially Holger Angenent and Sebastian Potthoff. We are grateful to Prof. Dr. Stefan Raunser for support and providing us access to the cryo-EM infrastructure of the Max Planck Institute (MPI) for Molecular Physiology Dortmund, Germany. We thank Dr. O. Hofnagel and Dr. D. Prumbaum (Dortmund) for assistance with cryo-EM dataset acquisition. We thank Dr. Alexander Belyy (Max-Planck Institute Dortmund, Germany) for kindly providing us *S. cerevisiae* MH272/3fα wild type strain. C.G. is a member of the Cells-in-Motion Interfaculty center (CiM) and a faculty member of CiM-IMPRS, a joint graduate program of the International Max Planck Research School (IMPRS) Molecular Biomedicine and the Graduate School of CiM at the University of Münster. M.R. is a member of CiM-IMPRS.

## Data Availability

The cryo-EM maps of “single-seam” and “twin-seam” state have been deposited to the Electron Microscopy Data Bank (EMDB) under the accession codes XXX and XXX. The cryo-EM dataset has been deposited to EMPIAR under accession codes XXX. The coordinates of the corresponding models have been deposited to the Protein Data Bank (PDB) under accession codes XXX and XXX. Other data are available from the corresponding author upon request.

## References

1. Tolbert, N. E. Microbodies-Peroxisomes and Glyoxysomes. Annu. Rev. Plant. Physiol. 22, 45–74; 10.1146/annurev.pp.22.060171.000401 (1971).

2. van den Bosch, H., Schutgens, R. B., Wanders, R. J. & Tager, J. M. Biochemistry of peroxisomes. Annual review of biochemistry 61, 157–197; 10.1146/annurev.bi.61.070192.001105 (1992).

3. Imanaka, T. & Shimozawa, N. Peroxisomes: Biogenesis, Function, and Role in Human Disease; 10.1007/978-981-15-1169-1 (2019).

4. Waterham, H. R. & Ebberink, M. S. Genetics and molecular basis of human peroxisome biogenesis disorders. Biochimica et biophysica acta 1822, 1430–1441; 10.1016/j.bbadis.2012.04.006 (2012).

5. Wanders, R. J. A. & Waterham, H. R. Peroxisomal disorders I: biochemistry and genetics of peroxisome biogenesis disorders. Clinical genetics 67, 107–133; 10.1111/j.1399-0004.2004.00329.x (2005).

6. Honsho, M., Okumoto, K., Tamura, S. & Fujiki, Y. Peroxisome Biogenesis Disorders. Advances in experimental medicine and biology 1299, 45–54; 10.1007/978-3-030-60204-8_4 (2020).

7. Dahabieh, M. S. et al. Peroxisomes and cancer: The role of a metabolic specialist in a disease of aberrant metabolism. Biochimica et biophysica acta. Reviews on cancer 1870, 103–121; 10.1016/j.bbcan.2018.07.004 (2018).

8. Fransen, M., Nordgren, M., Wang, B., Apanasets, O. & van Veldhoven, P. P. Aging, age-related diseases and peroxisomes. Sub-cellular biochemistry 69, 45–65; 10.1007/978-94-007-6889-5_3 (2013).

9. Cipolla, C. M. & Lodhi, I. J. Peroxisomal Dysfunction in Age-Related Diseases. Trends in endocrinology and metabolism: TEM 28, 297–308; 10.1016/j.tem.2016.12.003 (2017).

10. McNew, J. A. & Goodman, J. M. An oligomeric protein is imported into peroxisomes in vivo. The Journal of cell biology 127, 1245–1257; 10.1083/jcb.127.5.1245 (1994).

11. Gould, S. J., Keller, G. A., Hosken, N., Wilkinson, J. & Subramani, S. A conserved tripeptide sorts proteins to peroxisomes. The Journal of cell biology 108, 1657–1664; 10.1083/jcb.108.5.1657 (1989).

12. Lill, P. et al. Towards the molecular architecture of the peroxisomal receptor docking complex. Proceedings of the National Academy of Sciences of the United States of America, 33216–33224; 10.1073/pnas.2009502117 (2020).

13. Rüttermann, M. & Gatsogiannis, C. Good things come to those who bait: the peroxisomal docking complex. Biological chemistry 0; 10.1515/hsz-2022-0161 (2022).

14. Meinecke, M. et al. The peroxisomal importomer constitutes a large and highly dynamic pore. Nature cell biology 12, 273–277; 10.1038/ncb2027 (2010).

15. Dias, A. F. et al. The peroxisomal matrix protein translocon is a large cavity-forming protein assembly into which PEX5 protein enters to release its cargo. The Journal of biological chemistry 292, 15287–15300; 10.1074/jbc.M117.805044 (2017).

16. Skowyra, M. L. & Rapoport, T. A. PEX5 translocation into and out of peroxisomes drives matrix protein import. Molecular cell, 3209-3225.e7; 10.1016/j.molcel.2022.07.004 (2022).

17. Erdmann, R. & Schliebs, W. Peroxisomal matrix protein import: the transient pore model. Nature reviews. Molecular cell biology 6, 738–742; 10.1038/nrm1710 (2005).

18. Platta, H. W. et al. Ubiquitination of the peroxisomal import receptor Pex5p is required for its recycling. The Journal of cell biology 177, 197–204; 10.1083/jcb.200611012 (2007).

19. Feng, P. et al. A peroxisomal ubiquitin ligase complex forms a retrotranslocation channel. Nature 607, 374–380; 10.1038/s41586-022-04903-x (2022).

20. Sargent, G. et al. PEX2 is the E3 ubiquitin ligase required for pexophagy during starvation. The Journal of cell biology 214, 677–690; 10.1083/jcb.201511034 (2016).

21. Birschmann, I. et al. Pex15p of Saccharomyces cerevisiae provides a molecular basis for recruitment of the AAA peroxin Pex6p to peroxisomal membranes. Molecular biology of the cell 14, 2226–2236; 10.1091/mbc.e02-11-0752 (2003).

22. Matsumoto, N., Tamura, S. & Fujiki, Y. The pathogenic peroxin Pex26p recruits the Pex1p-Pex6p AAA ATPase complexes to peroxisomes. Nature cell biology 5, 454–460; 10.1038/ncb982 (2003).

23. Gardner, B. M. et al. The peroxisomal AAA-ATPase Pex1/Pex6 unfolds substrates by processive threading. Nature communications 9, 135; 10.1038/s41467-017-02474-4 (2018).

24. Pedrosa, A. G. et al. Peroxisomal monoubiquitinated PEX5 interacts with the AAA ATPases PEX1 and PEX6 and is unfolded during its dislocation into the cytosol. The Journal of biological chemistry 293, 11553–11563; 10.1074/jbc.RA118.003669 (2018).

25. Miyata, N. & Fujiki, Y. Shuttling mechanism of peroxisome targeting signal type 1 receptor Pex5: ATP-independent import and ATP-dependent export. Molecular and cellular biology 25, 10822–10832; 10.1128/MCB.25.24.10822-10832.2005 (2005).

26. Francisco, T. et al. A cargo-centered perspective on the PEX5 receptor-mediated peroxisomal protein import pathway. The Journal of biological chemistry 288, 29151–29159; 10.1074/jbc.M113.487140 (2013).

27. Rosenkranz, K. et al. Functional association of the AAA complex and the peroxisomal importomer. The FEBS journal 273, 3804–3815; 10.1111/j.1742-4658.2006.05388.x (2006).

28. Platta, H. W. et al. Regulation of peroxisomal matrix protein import by ubiquitination. Biochimica et biophysica acta 1863, 838–849; 10.1016/j.bbamcr.2015.09.010 (2016).

29. Schwerter, D. P., Grimm, I., Platta, H. W. & Erdmann, R. ATP-driven processes of peroxisomal matrix protein import. Biological chemistry 398, 607–624; 10.1515/hsz-2016-0293 (2017).

30. Erdmann, R. et al. PAS1, a yeast gene required for peroxisome biogenesis, encodes a member of a novel family of putative ATPases. Cell 64, 499–510; 10.1016/0092-8674(91)90234-p (1991).

31. Schliebs, W., Girzalsky, W. & Erdmann, R. Peroxisomal protein import and ERAD: variations on a common theme. Nature reviews. Molecular cell biology 11, 885–890; 10.1038/nrm3008 (2010).

32. Yu, H., Kamber, R. A. & Denic, V. The peroxisomal exportomer directly inhibits phosphoactivation of the pexophagy receptor Atg36 to suppress pexophagy in yeast. eLife 11, e74531; 10.7554/eLife.74531 (2022).

33. Law, K. B. et al. The peroxisomal AAA ATPase complex prevents pexophagy and development of peroxisome biogenesis disorders. Autophagy 13, 868–884; 10.1080/15548627.2017.1291470 (2017).

34. Ciniawsky, S. et al. Molecular snapshots of the Pex1/6 AAA+ complex in action. Nature communications 6, 7331; 10.1038/ncomms8331 (2015).

35. Meyer, H. & Weihl, C. C. The VCP/p97 system at a glance: connecting cellular function to disease pathogenesis. Journal of cell science 127, 3877–3883; 10.1242/jcs.093831 (2014).

36. Blok, N. B. et al. Unique double-ring structure of the peroxisomal Pex1/Pex6 ATPase complex revealed by cryo-electron microscopy. Proceedings of the National Academy of Sciences of the United States of America 112, E4017–25; 10.1073/pnas.1500257112 (2015).

37. Gardner, B. M., Chowdhury, S., Lander, G. C. & Martin, A. The Pex1/Pex6 complex is a heterohexameric AAA+ motor with alternating and highly coordinated subunits. Journal of molecular biology 427, 1375–1388; 10.1016/j.jmb.2015.01.019 (2015).

38. Saffian, D., Grimm, I., Girzalsky, W. & Erdmann, R. ATP-dependent assembly of the heteromeric Pex1p-Pex6p-complex of the peroxisomal matrix protein import machinery. Journal of structural biology 179, 126–132; 10.1016/j.jsb.2012.06.002 (2012).

39. Schwerter, D., Grimm, I., Girzalsky, W. & Erdmann, R. Receptor recognition by the peroxisomal AAA complex depends on the presence of the ubiquitin moiety and is mediated by Pex1p. The Journal of biological chemistry 293, 15458–15470; 10.1074/jbc.RA118.003936 (2018).

40. Debelyy, M. O. et al. Ubp15p, a ubiquitin hydrolase associated with the peroxisomal export machinery. The Journal of biological chemistry 286, 28223–28234; 10.1074/jbc.M111.238600 (2011).

41. Judy, R. M., Sheedy, C. J. & Gardner, B. M. Insights into the Structure and Function of the Pex1/Pex6 AAA-ATPase in Peroxisome Homeostasis. Cells 11; 10.3390/cells11132067 (2022).

42. Pan, M. et al. Mechanistic insight into substrate processing and allosteric inhibition of human p97. Nature structural & molecular biology 28, 614–625; 10.1038/s41594-021-00617-2 (2021).

43. Cooney, I. et al. Structure of the Cdc48 segregase in the act of unfolding an authentic substrate. Science (New York, N.Y.) 365, 502–505; 10.1126/science.aax0486 (2019).

44. L. Peña, A. H. de, Goodall, E. A., Gates, S. N., Lander, G. C. & Martin, A. Substrate-engaged 26S proteasome structures reveal mechanisms for ATP-hydrolysis-driven translocation. Science (New York, N.Y.) 362, eaav0725; 10.1126/science.aav0725 (2018).

45. Twomey, E. C. et al. Substrate processing by the Cdc48 ATPase complex is initiated by ubiquitin unfolding. Science (New York, N.Y.) 365, eaax1033; 10.1126/science.aax1033 (2019).

46. Wald, J. et al. Mechanism of AAA+ ATPase-mediated RuvAB-Holliday junction branch migration. Nature 609, 630–639; 10.1038/s41586-022-05121-1 (2022).

47. Lo, Y.-H. et al. Cryo-EM structure of the essential ribosome assembly AAA-ATPase Rix7. Nature communications 10, 513; 10.1038/s41467-019-08373-0 (2019).

48. Puchades, C. et al. Structure of the mitochondrial inner membrane AAA+ protease YME1 gives insight into substrate processing. Science (New York, N.Y.) 358, eaao0464; 10.1126/science.aao0464 (2017).

49. Weibezahn, J., Schlieker, C., Bukau, B. & Mogk, A. Characterization of a trap mutant of the AAA+ chaperone ClpB. The Journal of biological chemistry 278, 32608–32617; 10.1074/jbc.M303653200 (2003).

50. Hanson, P. I. & Whiteheart, S. W. AAA+ proteins: have engine, will work. Nature reviews. Molecular cell biology 6, 519–529; 10.1038/nrm1684 (2005).

51. Huang, R., Ripstein, Z. A., Rubinstein, J. L. & Kay, L. E. Cooperative subunit dynamics modulate p97 function. Proceedings of the National Academy of Sciences of the United States of America 116, 158–167; 10.1073/pnas.1815495116 (2019).

52. Bulfer, S. L., Chou, T.-F. & Arkin, M. R. p97 Disease Mutations Modulate Nucleotide-Induced Conformation to Alter Protein-Protein Interactions. ACS chemical biology 11, 2112–2116; 10.1021/acschembio.6b00350 (2016).

53. Banerjee, S. et al. 2.3 Å resolution cryo-EM structure of human p97 and mechanism of allosteric inhibition. Science (New York, N.Y.) 351, 871–875; 10.1126/science.aad7974 (2016).

54. Rao, M. V., Williams, D. R., Cocklin, S. & Loll, P. J. Interaction between the AAA+ ATPase p97 and its cofactor ataxin3 in health and disease: Nucleotide-induced conformational changes regulate cofactor binding. The Journal of biological chemistry 292, 18392–18407; 10.1074/jbc.M117.806281 (2017).

55. Puchades, C., Sandate, C. R. & Lander, G. C. The molecular principles governing the activity and functional diversity of AAA+ proteins. Nature reviews. Molecular cell biology 21, 43–58; 10.1038/s41580-019-0183-6 (2020).

56. Gates, S. N. & Martin, A. Stairway to translocation: AAA+ motor structures reveal the mechanisms of ATP-dependent substrate translocation. Protein science : a publication of the Protein Society 29, 407–419; 10.1002/pro.3743 (2020).

57. Li, S. et al. Molecular basis for ATPase-powered substrate translocation by the Lon AAA+ protease. The Journal of biological chemistry 297, 101239; 10.1016/j.jbc.2021.101239 (2021).

58. Wehmer, M. et al. Structural insights into the functional cycle of the ATPase module of the 26S proteasome. Proceedings of the National Academy of Sciences of the United States of America 114, 1305–1310; 10.1073/pnas.1621129114 (2017).

59. Wang, L., Myasnikov, A., Pan, X. & Walter, P. Structure of the AAA protein Msp1 reveals mechanism of mislocalized membrane protein extraction. eLife 9, e54031; 10.7554/eLife.54031 (2020).

60. Schieferdecker, A. & Wendler, P. Structural Mapping of Missense Mutations in the Pex1/Pex6 Complex. International journal of molecular sciences 20, 3756; 10.3390/ijms20153756 (2019).

61. Tan, D., Blok, N. B., Rapoport, T. A. & Walz, T. Structures of the double-ring AAA ATPase Pex1-Pex6 involved in peroxisome biogenesis. The FEBS journal 283, 986–992; 10.1111/febs.13569 (2016).

62. Grimm, I., Saffian, D., Girzalsky, W. & Erdmann, R. Nucleotide-dependent assembly of the peroxisomal receptor export complex. Scientific reports 6, 19838; 10.1038/srep19838 (2016).

63. Huang, R., Ripstein, Z. A., Rubinstein, J. L. & Kay, L. E. Probing Cooperativity of N-Terminal Domain Orientations in the p97 Molecular Machine: Synergy Between NMR Spectroscopy and Cryo-EM. Angewandte Chemie (International ed. in English) 59, 22423–22426; 10.1002/anie.202009767 (2020).

64. Shiozawa, K. et al. Structure of the N-terminal domain of PEX1 AAA-ATPase. Characterization of a putative adaptor-binding domain. The Journal of biological chemistry 279, 50060–50068; 10.1074/jbc.M407837200 (2004).

65. Shiozawa, K. et al. The common phospholipid-binding activity of the N-terminal domains of PEX1 and VCP/p97. The FEBS journal 273, 4959–4971; 10.1111/j.1742-4658.2006.05494.x (2006).

66. Birschmann, I., Rosenkranz, K., Erdmann, R. & Kunau, W.-H. Structural and functional analysis of the interaction of the AAA-peroxins Pex1p and Pex6p. The FEBS journal 272, 47–58; 10.1111/j.1432-1033.2004.04393.x (2005).

67. Agne, B. et al. Pex8p: An Intraperoxisomal Organizer of the Peroxisomal Import Machinery. Molecular cell 11, 635–646; 10.1016/S1097-2765(03)00062-5 (2003).

