## Supplementary Material for "Structure of the peroxisomal Pex1/Pex6 ATPase complex bound to a substrate"

*To whom correspondence should be addressed.

### Material and Methods

#### Cloning

*S. cerevisiae* Pex6 (NCBI gene number 855387 containing the E832Q mutation) and Pex1 (NCBI gene number 853636, WT) were cloned with a Gibson assembly into a pESC-URA vector (Agilent technologies, Santa Clara, USA). An N-terminal His-tag and a C-terminal $strep_2$-tag has been cloned to Pex6(E832Q) and Pex1, respectively. All constructs were verified by sequencing of the inserts.

#### Protein expression and purification

The generated plasmid was transformed via the lithium acetate method into MH 272/3fα *S. cerevisiae* cells[1] and successfully transformed cells were selected on SD media plates lacking uracil. One colony has been picked and inoculated in SD liquid media for preculture at 30°C for 12 h. The preculture has been shifted to fresh SD media and incubated at 30°C for 24 h. Co-expression was induced upon addition of 1/3 volume YEPG containing 2% (w/v) D-galactose. The expression has been further incubated at 30°C for 16-17 h. Cells were sedimented at 5,000 rpm for 10 min. and 4°C in an Avanti JC20XP (Beckman coulter, Brea, USA) centrifuge equipped with a JLA 8.100 (Beckman coulter, Brea, USA) rotor. The 35 g cell pellet was washed with deionized $H_2O$ and either further processed or frozen at -80°C.

For lysis, the cells were resuspended in lysis buffer (50 mM TRIS, pH 8, 300 mM NaCl, 5 mM $MgCl_2$, 1 mM ATP, 1 mM TCEP, 5 % Glycerol, 1x Protease Inhibitor) and drop-wise frozen in liquid nitrogen to form small beads. The frozen cells were lysed in a ZM 200 ultra-centrifugal mill (Retsch GmbH, Haan, Germany) precooled with liquid nitrogen at 18,500 rpm until a homogeneous powder was obtained.

The cell lysate was thawed, and debris were removed by centrifugation at 25,000 rpm (SA25.50 rotor, Beckman Coulter, Brea USA) for 45 minutes at 4°C using an Optima XPN-80 (Beckman Coulter, USA). The supernatant was subsequently further cleared by an additional ultracentrifugation step at 40,000 rpm (SA25.50 rotor, Beckman

Coulter, Brea, USA) for 15 minutes at 4°C and filtration with a 0.45 µm nitrocellulose filter.

For strep-tactin purification, 300 ml filtered supernatant were subjected to 1.5 ml strep-tactin super flow high-capacity beads (IBA Lifesciences GmbH, Göttingen, Germany) in a gravity flow column. The beads were washed with 10 column volumes strep washing buffer (50 mM TRIS, pH 7.4, 300 mM NaCl, 3mM ATP, 5 mM $MgCl_2$, 1 mM TCEP, 5 % Glycerol). The protein was eluted with strep elution buffer (50 mM TRIS, pH 7.4, 300 mM NaCl, 1 mM ATP, 5 mM $MgCl_2$, 1 mM TCEP, 5 % Glycerol, 10 mM desthiobiotin). Collected fractions containing the proteins of interest were concentrated using a 100 kDa MWCO centrifugal filter at 2,500 rpm and 4°C.

The concentrated protein fractions were separated by size on a Superose 6 10/300 column operated with an AEKTA Purifier (GE Healthcare Bioscience, Chicago, USA). UV-absorption was monitored and Pex1/Pex6(E832Q) fractions were pooled and concentrated.

Pooled elutes were concentrated to 400 µl using a MWCO 100kDa. The concentrated sample was cleared from aggregates by centrifugation at 10,000 rpm and 4°C. The supernatant was further purified via size-exclusion chromatography on a Superose 6 Increase 10/300 column (GE Healthcare Bioscience, Chicago, USA). Peak fractions were pooled and concentrated to a final protein concentration of 3.6 mg/ml.

To identify fractions of interest, Mini-PROTEAN TGX precast gels (Bio-Rad, USA) were used as recommended by the manufacturer. Protein bands were visualized with UV illumination or semi-dry blotted to Trans-Blot Turbo Transfer Packs (Bio-Rad, Hercules, USA). Immunolabeling was performed with horse-reddish peroxidase (HRP) coupled strep-tactin or anti-His antibodies.

**Negative stain electron microscopy**

Carbon coated copper grids (Agar scientific, Stansted Mountfitchet, United Kingdom; G2400C) were glow discharged as described previously[2] and then treated with Poly-

L-lysine by incubating a droplet of 3 µl 0.1 % (w/v) poly-L-lysine hydrobromide (Sigma-Aldrich, St. Louis, USA) for 30 seconds on the grid. Excess fluid was subsequently blotted with Whatman paper (No 5.), and the grids were washed two times with 10 µl water. After the grids were dried, 4µl of purified Pex1-strep$_2$/His-Pex6(E832Q) complex was applied for 2 minutes at room temperature. The sample was blotted, washed with 10 µl 0.75% uranylformate solution, and blotted again. Another 10 µl 0.75% uranylformate solution was applied to the grid and incubated for 45 seconds at RT.

Images were recorded with a JEM-1400 (JEOL) equipped with LaB6 cathode and a 4K CMOS detector F416 (TVIPS). Images were acquired at a pixel size of 1.84 Å.

#### Cryo-EM sample preparation and data acquisition

For cryo-EM sample preparation, 4 µl protein solution (1.8 mg/ml) were applied to a glow-discharged holey carbon grid (R 2/1, 200 mesh, Quantifoil Micro Tools GmbH, Großlöbichau, GER). Grids were blotted for 3.5 sec and plunge-frozen after 1 sec drain time at 100% humidity in liquid ethane using a Vitrobot II (FEI, Hillsboro, USA). The plunged EM grids were used for automatically recording of 16,763 movies using the EPU software with a Titan Krios (FEI, Hillsboro, USA) with Cs-corrector and XEFG electron source at 300 kV. The cryo-EM data collection parameters are summarized in Supplementary Table 1.

#### CryoEM data processing

The resolutions of cryo-EM maps are given according to the gold standard Fourier shell correlation (FSC) after combination of the half maps applying a full particle mask unless otherwise stated. During dataset acquisition, the quality of the data was monitored and the data was transferred using TranSPHIRE[3]. The super resolution movies (0.34 Å/pixel) were subjected to motion correction using Motioncor2 1.3.2 at 0.68 Å/pixel[4] and the Contrast Transfer Function (CTF) was estimated using CTFFIND 4.1.14 [5]. Particles in representative micrographs were manually picked to train a model, which was then used for automated particle picking in crYOLO 1.8.1[6], selecting

1,259,079 particles with a box size of 384 x 384 pixels. Subsequently, 2D classification was performed using the iterative stable alignment clustering method (ISAC) with 200 particles/class[7,8]. After manually sorting out the "low quality" 2D classes, 605,359 particles remained. A 3D reconstruction (4.5 Å) was computed with MERIDIEN[9] using a negative stain EM structure (EMDB-2585) as reference[10]. The particles were further processed in RELION 3.1.4[11,12] using the reconstruction previously obtained with MERIDIEN. The particle overall demonstrated a pseudo C3-symmetry with more flexible N-terminal domains and an unsymmetric D2-ring. To improve overall resolution and avoid symmetry conflicts or particle misalignment during global 3D classification, particles were triplicated and given as an input for a 3D refinement imposing C3-symmetry reaching 4.5 Å. Subsequently, the C3-refined map was used as a template in 3D classification with C1-symmetry giving 6 classes. Then, duplicates have been removed from each individual 3D class. Finally, class 3 (147,925 particles) and class 4 (128,983 particles) representing two distinct conformations, were selected and further locally refined to an average resolution of 4.4 Å and 4.8 Å, respectively. Both classes were then improved with Bayesian polishing, per-particle CTF-refinement and another final round of Bayesian polishing to 4.1 Å (class 3) and 4.7 Å (class 4). The densities of class 3 and 4 were further improved with local anisotropic half map sharpening and density modification in Phenix 1.20[13,14]. The average resolution was improved to 3.9 Å (class 3) and 4.3 Å (class 4) resolution, respectively. Note that the resolution of density modified maps was determined via Phenix at FSC=0.5[14]. The overall processing workflow is shown in Supplementary Fig. 2. Class 3 ("single seam") has an average resolution of 3.7 Å for the D1 ring and 3.9 Å (FSC=0.143) for the D2 ring (improved to 3.3 Å (D1) and 3.6 Å (D2) after density modification) (Supplementary Fig. 3).

Class 4 shows an average resolution of 3.9 Å in the D1 and 4.7 Å in the D2 ring (improved to 3.8 (D1) to 4.3 (D2) Å after density modification) (Supplementary Fig. 10). Local resolution was estimated using SPHIRE and Phenix 1.20 for final reconstruction and density modified maps, respectively[9,13]. 3D FSC was computed using the 3DFSC server[15].

#### Model building of Pex1/Pex6

Molecular models of the Pex1/6 complex are not available in the PDB. Alphafold predictions[16] of the full-length *Saccharomyces cerevisiae* Pex1 and Pex6 subunits have been used for rigid body fitting into the best resolved cryo-EM map (class3) and the assignment of the N1, N2, D1 and D2 domains of Pex1 and Pex6.

For model building of class 3 ("single seam"), the Alphafold predictions of Pex1 and Pex6 were first split to individual domains (N1, N2, D1, D2). Note that Pex1(N1) was not resolved in both cryo-EM structures. The individual domains of Pex1 (N2,D1,D2) and Pex6 (N1,N2,D1,D2) were fit to the density-modified class 3 map using ChimeraX 1.4[17] and the model was built with Coot 0.9.8[18]. The less well-resolved N-terminal domains Pex1(N2) and Pex6(N1) (class 3: ~6-8 Å) were instead only flexibly fitted in the density using the iMODFit plugin in UCSF Chimera 1.15[19,20]. The better resolved D1 (class 3: 3.3 – 3.8 Å) and D2 (class 3: 3.4 – 3.8 Å, seam subunit: ~6 Å) domains were real space refined into the density modified maps using Phenix 1.20[13]. For the protrusion domain of Pex1 (helix α28) bound to the N2-terminal domain of Pex6, an Alphafold multimer model was predicted for this interface (Supplementary Fig. 9c) due to low local resolution (~8 Å). This model was rigid-body fitted into the density.

We observed a characteristic density in the channel of D2 rings corresponding to an unidentified endogenous substrate. We modeled a polyalanine peptide with an overall length of 10 residues. Densities at the putative nucleotide binding domains of both cryo-EM structures have been carefully analyzed. We fitted either ATP or ADP argued by the density, opening of the nucleotide pocket or position of R-fingers (Supplementary Fig. 5). $Mg^{2+}$ ions were placed and relaxed in such a way that they enter into the favorable coordination with the ATP as well as the threonine of the Walker A domain using the ChimeraX molecular dynamics plugin ISOLDE[21]. In a last step, sterically unfavorable conformations have been corrected using Coot 0.9.8 with iterative rounds of Phenix 1.20 real-space refinements in between[13]. The entire procedure was repeated to create an atomic model for the less well resolved class 4 cryo-EM density. Here we used the molecular model obtained for class 3 as a starting point. Refinement statistics are listed in Supplementary Table 1.

#### Additional Software

Images and movies were created using ChimeraX 1.4[17]. Figure 4b was prepared with the PyMol script modevectors with a cutoff value of 4.0 Å. The length of the arrows depicts the magnitude of the motion. Distance and surface area measurements have been done using ChimeraX 1.4. Sequence alignments have been performed with SnapGene 6.0.6. Interfaces have been analyzed with the PDBePISA server[22]. Secondary structures of Pex1 and Pex6 (Supplementary Fig. 4) have been computed with PDBsum[23].

**Supplementary Table 1:** Cryo-EM data collection and refinement statistics

| **Data collection** | |
|---|---|
| Microscope | Titan Krios with Cs-corrector and XFEG electron source |
| Voltage (kV) | 300 |
| Nominal magnification | 105,000 x |
| Electron Dose ($e^-$/ $Å^2$) | 60 |
| Number of frames | 60 |
| Detector | Gatan K3 |
| Pixel size (Å) | 0.34 (super resolution mode) |
| Defocus range (µm) | -0.6 to -2.4 |
| Micrographs | 16,763 |
| **Reconstruction** | |
| Software | SPHIRE 1.4, RELION 3.1.4, Phenix 1.20 |

| | | |
|---|---|---|
| Total extracted particles | 1,259,079 | |
| Structure<br>Pex1/Pex6E832Q | Class 3<br>Single-‘seam’ conformation | Class 4<br>Twin-‘seam’ conformation |
| Particles | 164,610 | 128,983 |
| Symmetry | C1 | C1 |
| Final average resolution, gold standard FSC=0.143 | 4.1 Å | 4.7 Å |
| Final average resolution after density modification FSC=0.5 | 3.9 Å | 4.3 Å |
| **Model refinement** | | |
| Peptide chains | 7 | 7 |
| Residues | 5566 | 5565 |
| Ligands | ATP: 10, ADP: 2, $Mg^{2+}$: 10 | ATP: 10, ADP: 2, $Mg^{2+}$: 10 |
| RMSD Bond length (Å) | 0.004 | 0.003 |
| RMSD Bond angles (°) | 0.745 | 0.733 |
| Ramachandran outliers (%) | 0.05 | 0.16 |
| Ramachandran allowed (%) | 9.47 | 13.49 |
| Ramachandran favored (%) | 90.47 | 86.34 |
| Rotamer outliers (%) | 0.04 | 0.02 |
| MolProbity score | 2.15 | 2.31 |
| Clash score | 12.53 | 14.9 |
| EMRinger score | 1.53 | 0.95 |

### Supplementary Figures

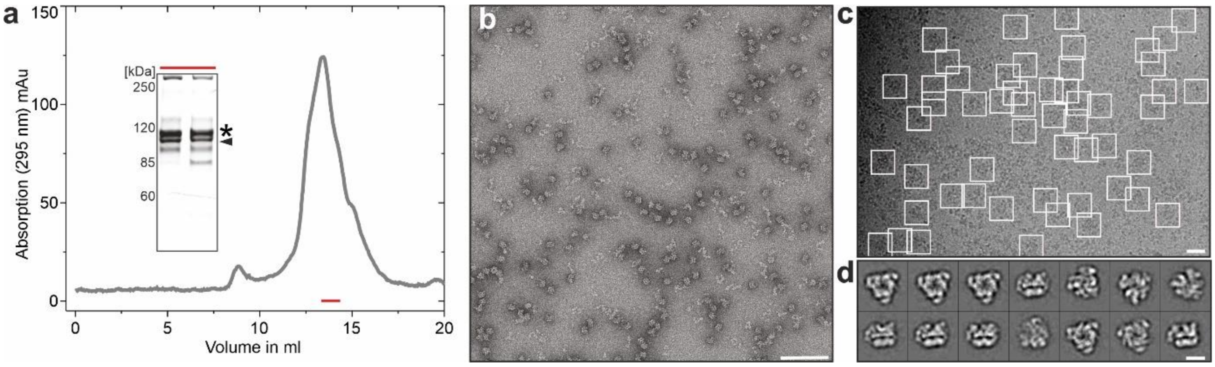


**Supplementary Figure 1: Expression and purification of recombinant Pex1/Pex6_WB. a)** Size exclusion chromatogram (SEC)**,** showing one major peak corresponding to a complex of Pex1 (star) and Pex6_WB (arrow), as confirmed by SDS-PAGE (inset). The red bar indicates fractions, which were pooled for subsequent cryo-EM studies. **b)** Representative micrograph of negatively stained Pex1/Pex6_WB complex after SEC. Scalebar 100 nm. **c)** Representative cryo-EM micrograph of Pex1/Pex6_WB low-pass filtered to 5 Å. Scale bar 20 nm. Boxes represent particles identified by crYOLO. **d)** Representative reference-free 2D class averages of Pex1/Pex6 Pex6_WB complex. Scale bar 10nm.

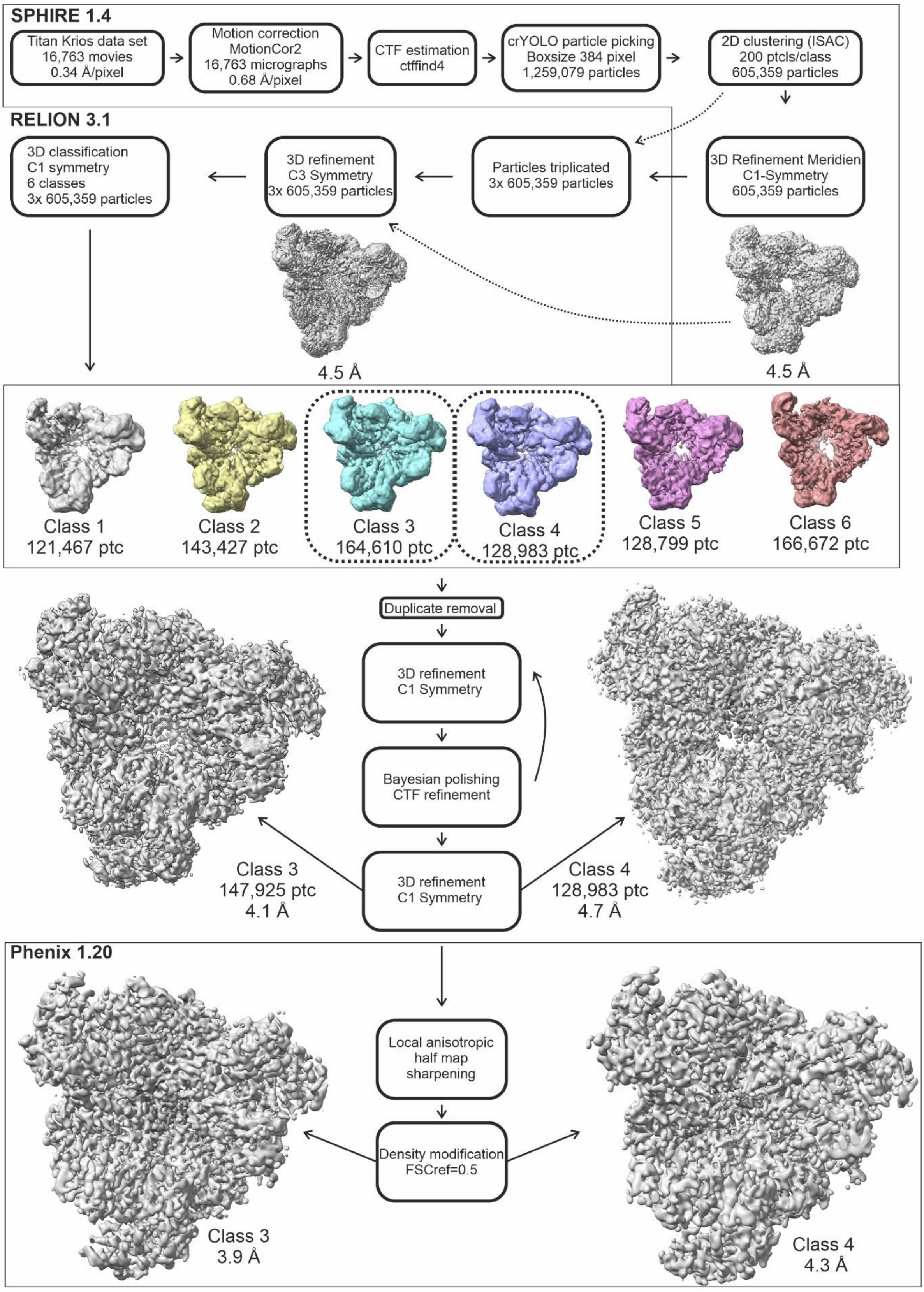


**Supplementary Figure 2: Single-particle cryo-EM processing workflow.** The final maps of class 3 and class 4 from RELION and Phenix have been used for model building.

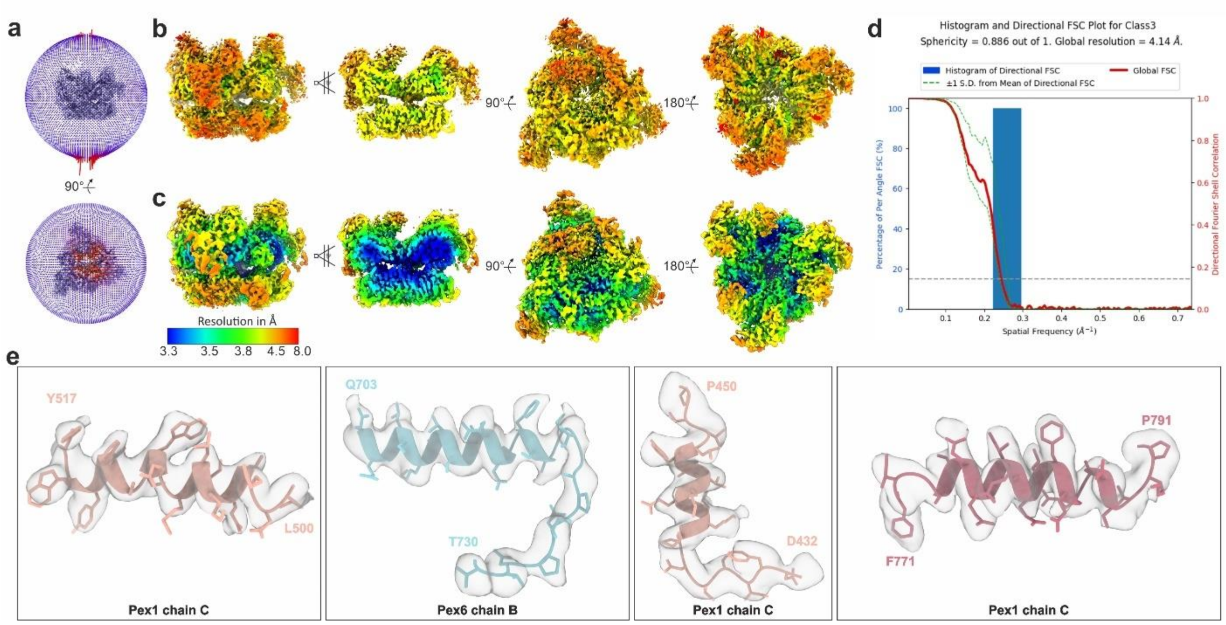


**Supplementary Figure 3: Cryo-EM structure of the single 'seam' state (class 3). a)** Angular distribution for the final round of the refinement. **b-c)** Different surface views and cross-section of the cryo-EM density map colored according to the local resolution, upon post-processing in RELION (b) and after local anisotropic half map sharpening and density modification using Phenix (c). **d)** 3D FSC curve. **e)** Representative areas of the density map superimposed with the molecular model.

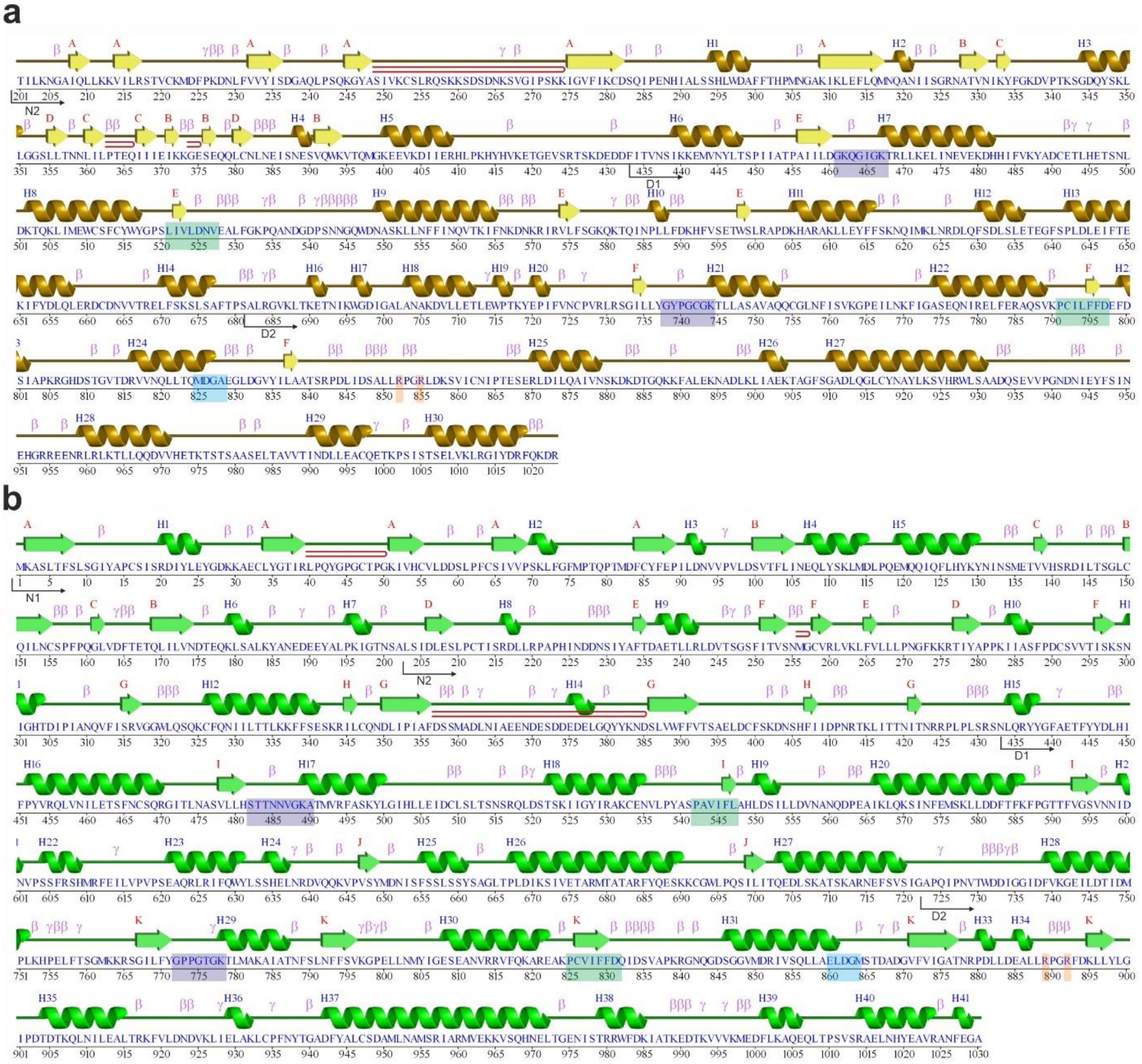

**Supplementary Figure 4. Secondary structure of Pex1(chain c) (a) and Pex6(chain d) (b)** computed with PDBsum (http://www.ebi.ac.uk/pdbsum/)[23]. α-helices are labeled H1, H2, …,H41 and β-strands by their sheets as A, B, …,K. Structural motifs β-turns, γ-turns, and β-hairpins are marked as β, γ, and ⊃, respectively. Important conserved motifs are indicated: Walker A (dark purple), Walker B (green), ISS (cyan), pore loops (red loop) and arginine fingers (orange).

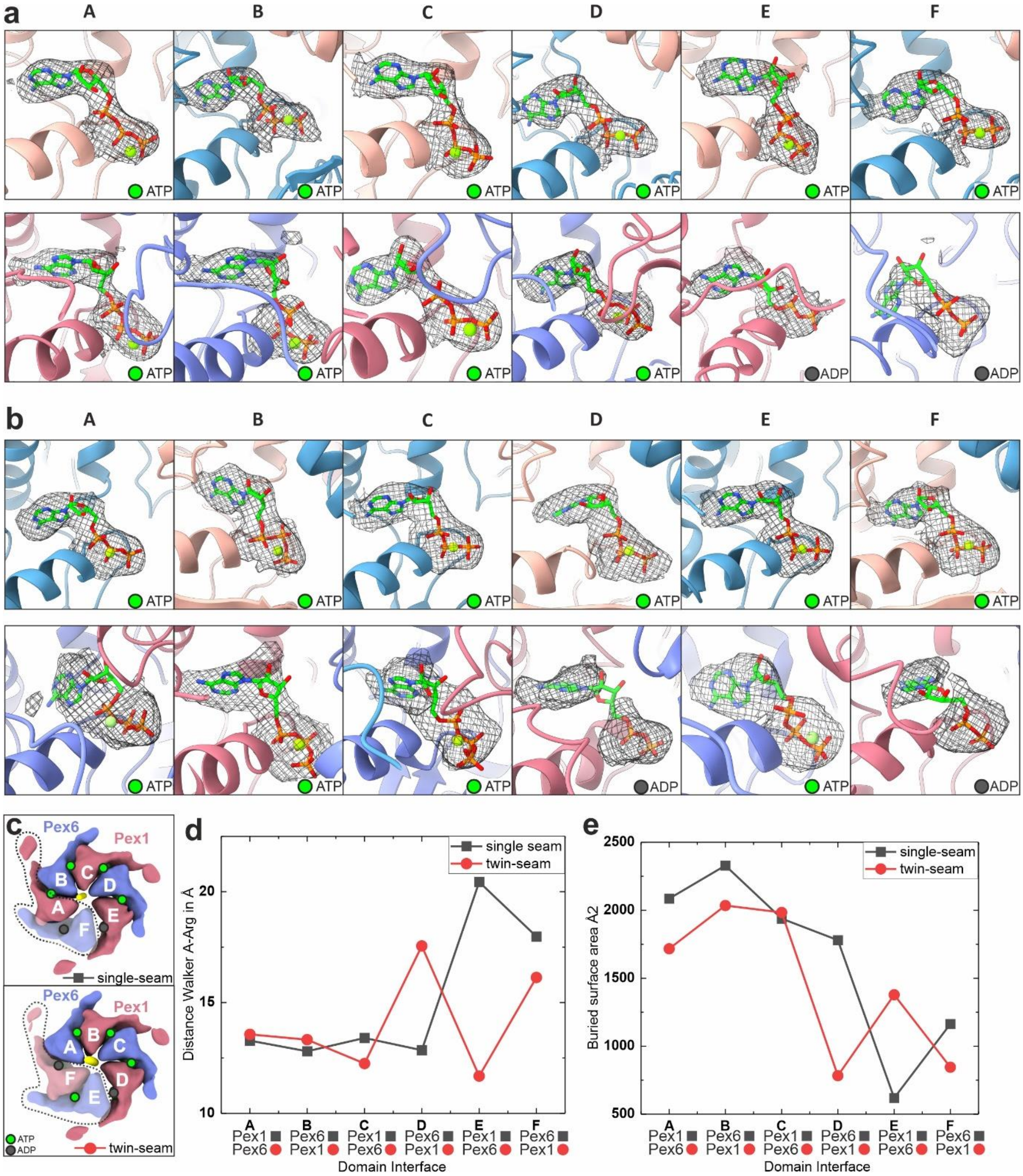


**Supplementary Figure 5: Analysis of the nucleotide binding pockets a-b)** Nucleotide models and corresponding cryo-EM density of the D1 (upper panels) and D2 (lower panels) ring of the single-seam (a) and twin-seam (b) state. **c)** Position of nucleotide binding pockets and D2-interfaces respective to the disengaged seam (F) domain for the single-seam (upper-inset) and twin-seam state (lower-inset). Letters (from A-F) define the individual subunit position within the stair-case. Domain A occupies the highest position of the spiral. **d)** Measurements of the opening of the nucleotide binding pocket for the single- (gray) and twin-seam (red) state. Shown are the distances between the Cα of Walker A Thr and the Cα of the Arg-finger of the neighboring subunit (Pex1-Pex6 interface: Pex1(T745)-Pex6 (R892); Pex6-Pex1 interface: Pex6(T779)-Pex1(R855). **e)** Contact area between the large and small AAA+ domains of neighboring D2 domains. The buried surface area was measured between the D2 domains of Pex1 (residues 681-1020) and Pex6 (residues 723-1030).

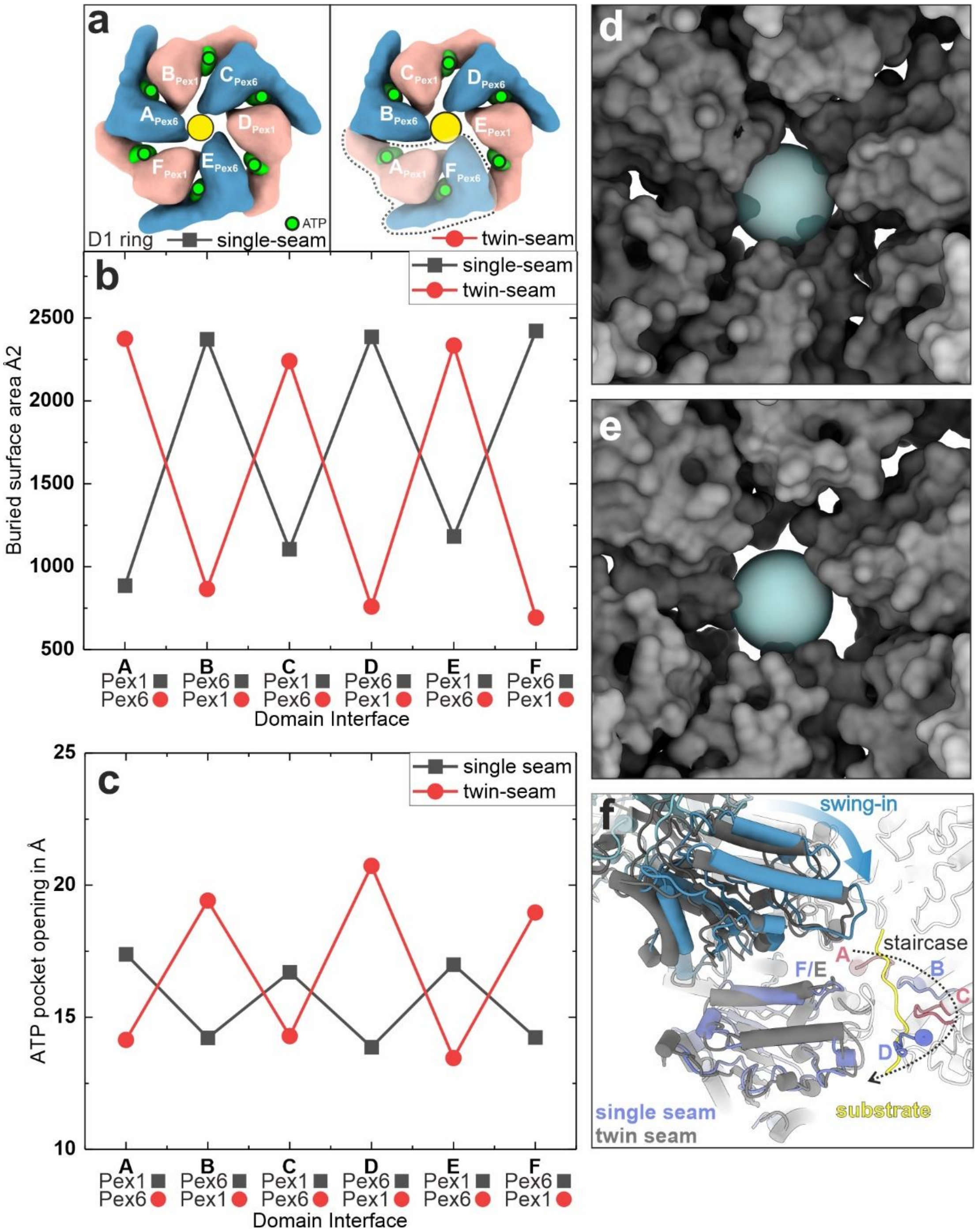


**Supplementary Figure 6: Structural dynamics of the D1 Ring between single- and twin-seam state. a)** Position of nucleotide binding pockets and D2-interfaces for the single-seam (left inset) and twin-seam state (right inset). **b-c)** Shown are measurements of the opening of the b) nucleotide binding pocket (Pex1 T468 to Pex6 D582 and Pex6 T491 to Pex1 K563) and the c) buried surface between adjacent subunits for the single- (gray) and twin-seam (red) state. The buried surface area was measured between the D2 domains of Pex1 (residues 201-680) and Pex6 (residues 201-722). **d-e)** D1 pore of d) single seam state and e) twin-seam state as surface representation. The blue transparent sphere has a diameter of 15 Å. Note the widening of the pore from single- to twin seam-state. **f)** Single-seam state (gray) superimposed on twin-seam state (colored) with chains D (except pore loop) removed to show the interior. Note that the twin-seam (left protomers, gray) swing in on the opposite side of the pore loops 1 staircase, during transition to single-seam (left protomer, colored).

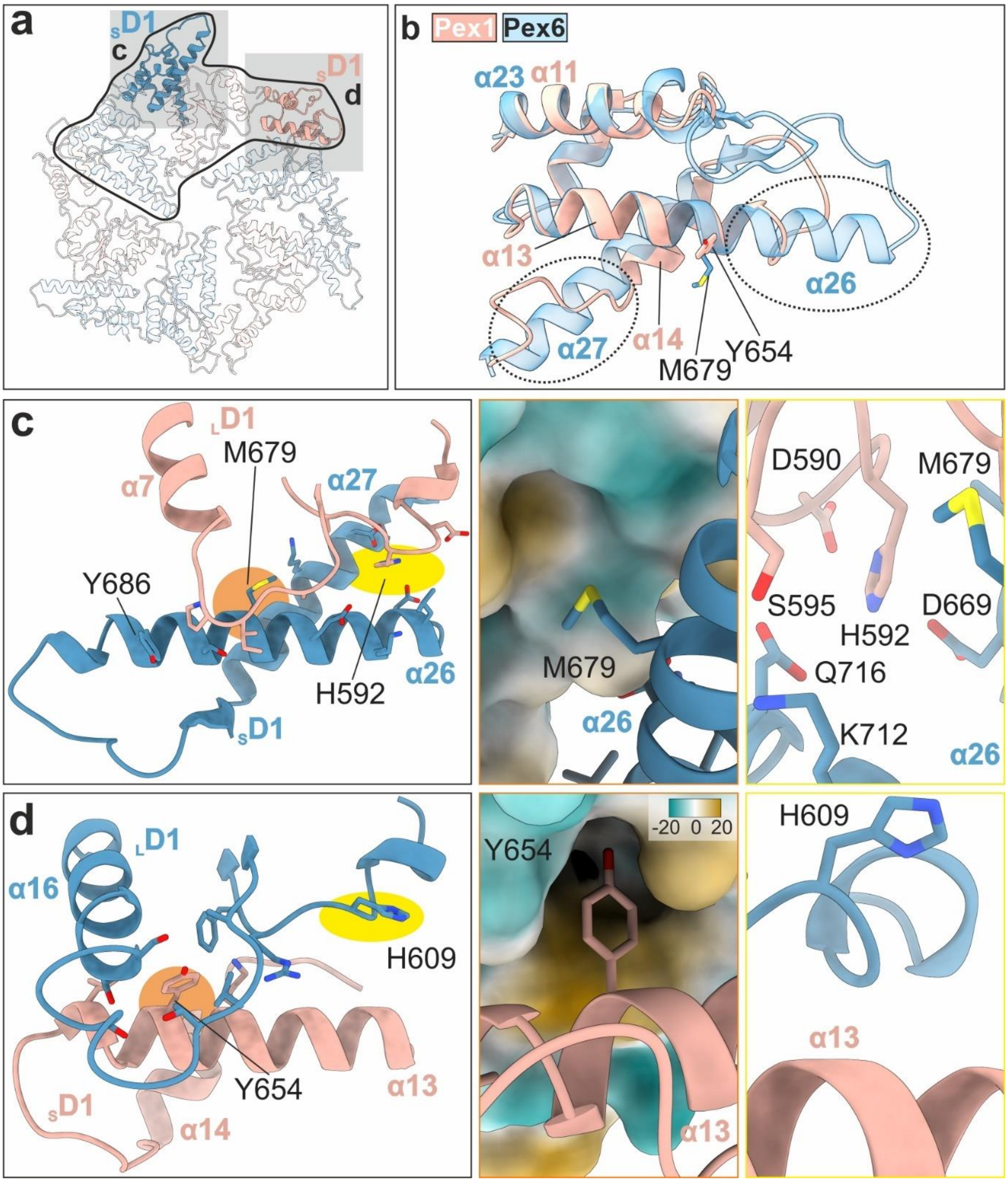


**Supplementary Figure 7: Distinct features of the small D1-ATPase domains of Pex1 and Pex6 determine the "trimer-of-dimers" arrangement in the D1 ring. a)** Top view of the Pex1/Pex6 D1 ring. The Pex6/Pex1 dimer indicated by a black line. The D1 small ATPase subdomains ($_S$D1) of Pex1 (beige) and Pex6 (blue) are highlighted. **b)** Structural alignment of Pex1($_S$D1) (residues 605-685, beige) and Pex6($_S$D1) (residues 621-721, blue transparent). Note the deletion in helix α13 and α14 of Pex1($_S$D1). **c)** Contact site of Pex6($_S$D1) with Pex1($_L$D1) (compact Interface within the dimer). M679 of the prolonged α26 helix of Pex6($_S$D1) (orange highlight) binds to a hydrophobic pocket formed by helix α7, and the loop upstream (residues 447-456) of Pex1($_L$D1). The middle panel shows a close-up view of this interface, with the surface of the pocket colored by hydrophobicity. The conserved histidine H592 of Pex1($_L$D1) (yellow highlight) binds into a negatively charged pocket formed by the prolonged helices α26 and α27 of Pex6($_S$D1). A close-up view of this interface is shown in the right panel.

**d)** Contact site of Pex1($_S$D1) with Pex6($_L$D1) (less compact interface between dimers). Due to the deletion in helix α13 of Pex1($_S$D1), Y654 (orange highlight) binds to the hydrophobic pocket formed by helix α16 and the loop upstream (residues 464-478) of Pex6($_L$D1). The middle panel shows a close-up view of this interface, with the surface of the pocket colored by hydrophobicity. Interestingly, due to the deletion, the analogous conserved H609 of Pex6 ($_L$D1) (yellow highlight, right panel) does not contact Pex1($_S$D1).

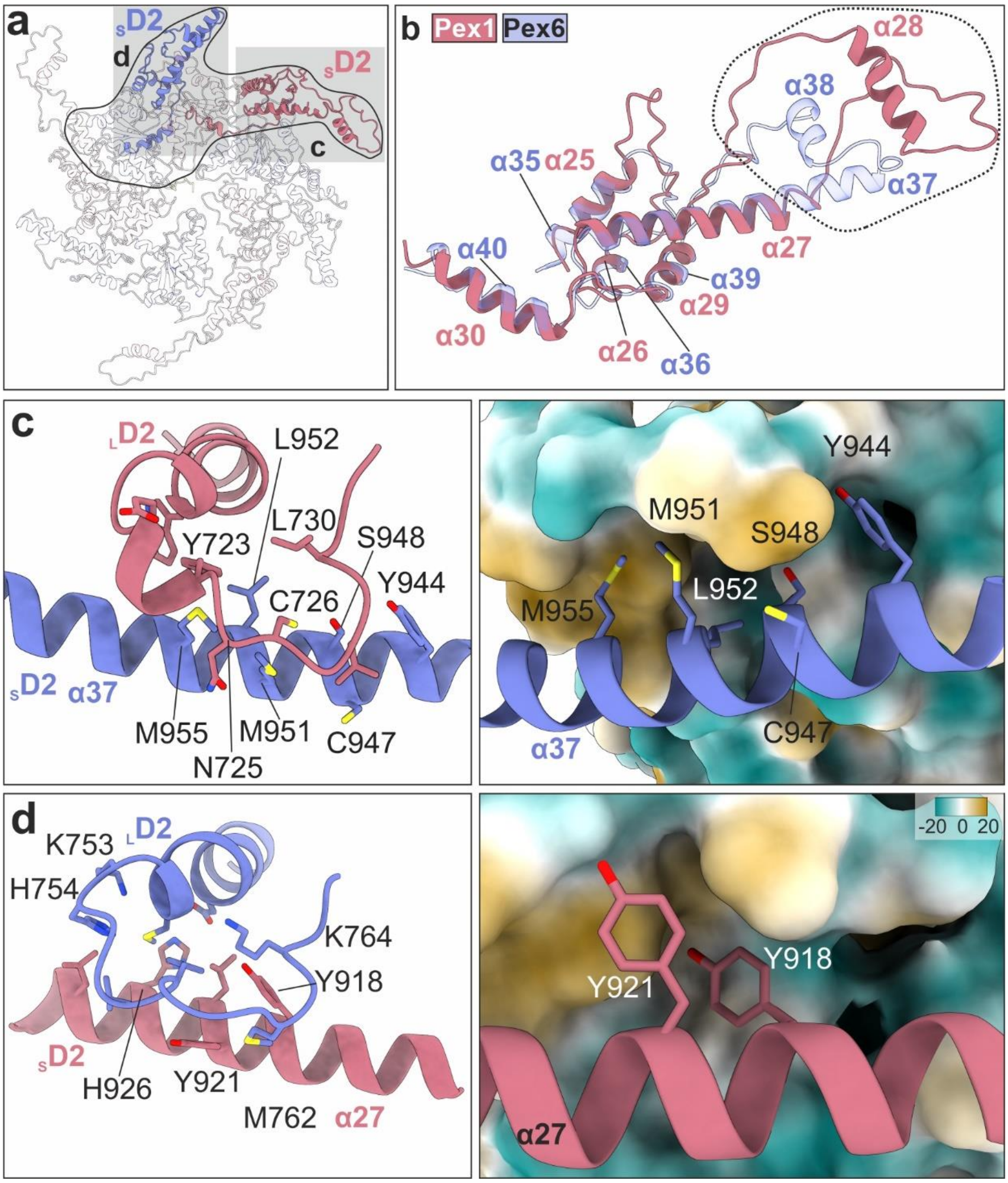


**Supplementary Figure 8: Distinct features of the small D2-ATPase domains fine-tune trimer-of-dimer arrangement in the D2-ring. a)** Top view of the Pex1/Pex6 D2 ring. The Pex6/Pex1 dimer is indicated by a black line. The D2 small ATPase subdomains ($_S$D2) and large ATPase domains ($_L$D2) of Pex1 (red) and Pex6 (blue) are highlighted. **b)** Structural alignment of Pex1($_S$D2) (residues 868-1021, beige) and Pex6($_S$D2) (residues 900-1030, blue transparent). Note the distinct Pex1(α28) and Pex6(α38) protrusion domains highlighted by the dashed line. **c)** Contact site of Pex6($_S$D2) with Pex1($_L$D2) (compact Interface within the dimer). Hydrophobic residues (M951, L952, M955 and Y944) mediate interactions within the dimer binding into a hydrophobic groove (right panel) formed by Pex1(α18) and the loop upstream.

**d)** Contact site of Pex1($_S$D2) with Pex6($_L$D2) (loose interface between dimers). Two hydrophobic residues (Y918 and Y921) on Pex1(α27) mediate interactions between dimers via binding into a hydrophobic pocket (right panel) formed by Pex6(α28) and the loop upstream.

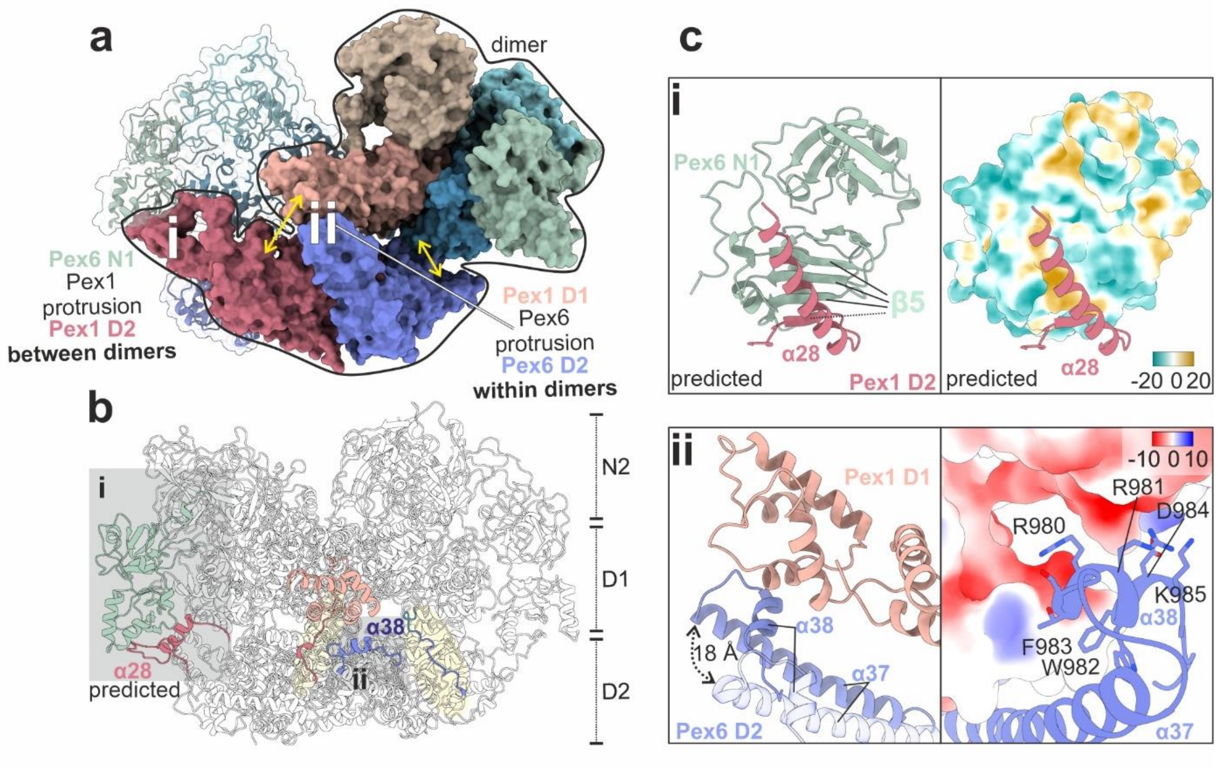


**Supplementary Figure 9: Interactions between the D1 and D2 ring. a)** Overview of the D1-D2 interactions via the protrusion domains of Pex1(i), Pex6(ii) and flexible linker peptides (yellow double-arrows). A Pex1/Pex6 subunit dimer is shown in surface representation. The clockwise (from top) adjacent Pex6 subunit is shown in ribbon representation. **b)** The structural elements involved in D1-D2 interactions are shown in ribbon representation and highlighted in color. In particular, the linker peptides (Pex1 (red, aa680-690); Pex6 (blue, residues 722-729)) covalently link the D1 and D2 ATPase cassettes. The protrusion domain of Pex1(D2) (amphipathic α28 helix) is flanked by long flexible loops and establishes an interface between neighboring dimers via anchoring at the adjacent Pex6(N1) domain (green) (i). The protrusion domain of Pex6(D2) (helix α38) mediates contact between the Pex6(D2) and the Pex1($_S$D1) domains within a subunit dimer (ii). **c)** Close-up view of interface (i) (upper-insets) and ii (lower insets). Please note: The Pex1(D2) protrusion domain (α28) is not well resolved. To address this, the interface (i) (Pex1(D2) (residues 940-976, red) and Pex6(N1)(1-201, green) (left panel) was predicted using Alphafold. Note that the Pex1 β-strand upstream of α28 enters the β-sheet 5 (β5) of Pex6. Together they form a hydrophobic groove, to which α28 (red) docks (right panel). **ii** This intra-dimer interface is flexible and depends on the nucleotide binding state of Pex6(D2). The Pex6(α38) protrusion helix at the tip of Pex6(α37) undergoes large movements (indicated by a dashed arrow) during ATPase cycle (right panel). During the transition from twin- to single seam state, α38 of the Pex6(D2) domain reengages and binds into a charged groove of Pex1(D1). Scalebar kcal/(mol·e) at 298 K.

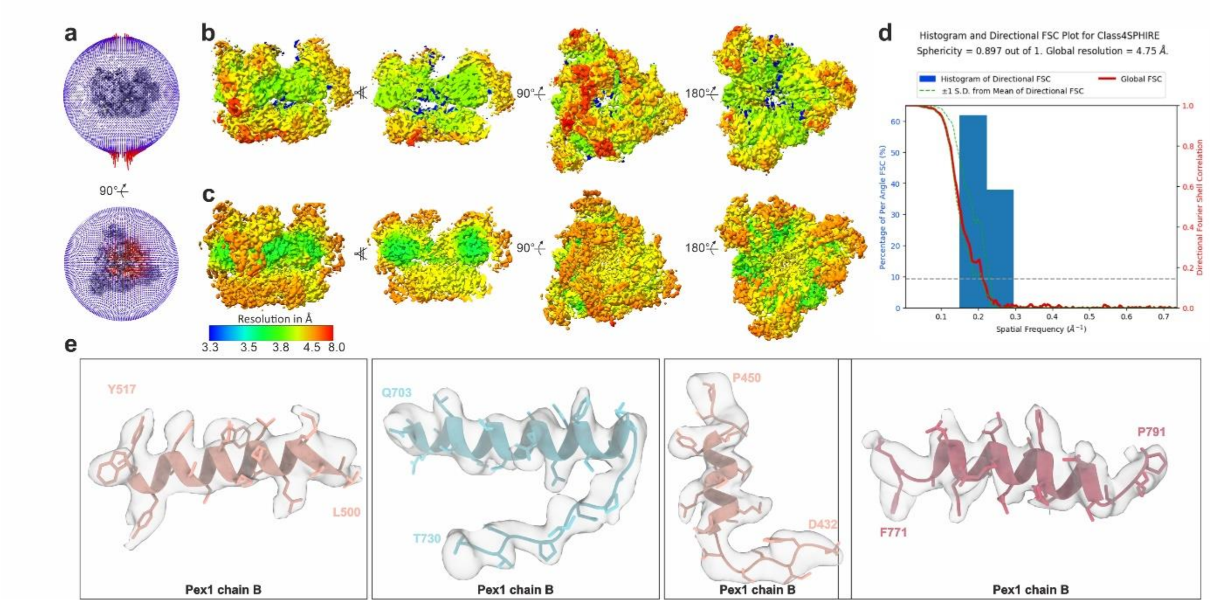


**Supplementary Figure 10: Cryo-EM structure of the single 'twin-seam' state (class 4). a)** Angular distribution for the final round of the refinement. **b-c)** Different surface views and cross-section of the cryo-EM density map colored according to the local resolution, upon post-processing in RELION (b) and after local anisotropic half map sharpening and density modification using Phenix (c). **d)** 3D FSC curve. **e)** Representative areas of the density map superimposed with the molecular model

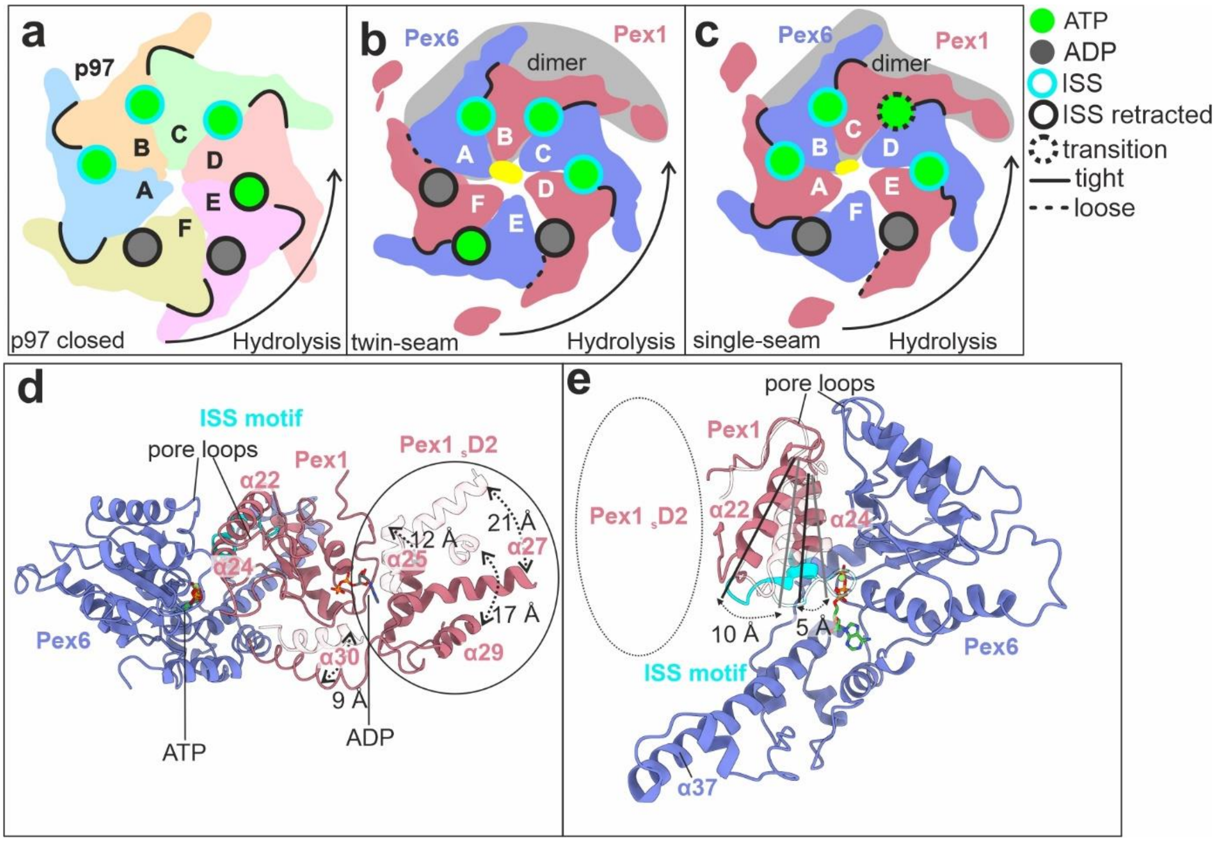


**Supplementary Figure 11: Inter-subunit signaling (ISS) motif controls ATP hydrolysis. a)** Illustration showing the arrangement of the D2 ring in p97 (closed conformation PDB: 7LN5)[24]. Note that the ISS motif of subunit F (ADP) is retracted from the nucleotide site of the neighboring subunit E and primes this site for hydrolysis. **b-c)** Illustration showing the arrangement of the D2 ring in Pex1/Pex6 in "single-seam" (b) and "twin-seam" state (c). Note that the ISS at the tight dimeric interface (D/E) ("single-seam", b) blocks hydrolysis. Upon release, the dimer (E/F) ("twin-seam", c) is less compact, the ISS is retracted and the dimeric interface is primed for hydrolysis. **d)** Pex6/Pex1 "twin-seam dimer" (chain E/F, colored blue/red) superimposed with staircase engaged Pex6/Pex1 dimer from the single-seam state (chain D/E, chain D removed, chain E transparent red) aligned on the large ATPase subdomain of Pex6 to visualize conformational changes upon dimer-detachment from the staircase. Note large conformational changes of the Pex1($_S$D2) subdomain upon detachment from the staircase. **e)** Same as in (**d**) but viewed from top. Pex1($_L$D2) is removing the ISS motif from the nucleotide pocket to prime it for hydrolysis.

### Supplementary Videos

**Supplementary Video 1 Cryo-EM structure of the Pex1/Pex6 (class 3; "single-seam" state).** Color code as in Figure 1a.

**Supplementary Video 2 Structural comparison between "twin-seam" and "single-seam" state.** Color code as in Figure 1a. Note that the transition from twin-seam to single-seam state in the D2 ring induces a swing-in of the Pex1(D1/N2)/Pex6(D1/N2/N1) domains in the D1 ring (see Fig. 4d-e).

**Supplementary Video 3 Structural comparison of the Pex1(D2)/Pex1(D1) interface between "twin-seam" and "single-seam" state.** (see Fig. 4g).

### References


1. Peisker, K. *et al.* Ribosome-associated complex binds to ribosomes in close proximity of Rpl31 at the exit of the polypeptide tunnel in yeast. *Molecular biology of the cell* **19,** 5279–5288; 10.1091/mbc.e08-06-0661 (2008).
2. Lill, P. *et al.* Towards the molecular architecture of the peroxisomal receptor docking complex. *Proceedings of the National Academy of Sciences of the United States of America,* 33216–33224; 10.1073/pnas.2009502117 (2020).
3. Stabrin, M. *et al.* TranSPHIRE: automated and feedback-optimized on-the-fly processing for cryo-EM. *Nature communications* **11,** 5716; 10.1038/s41467-020-19513-2. (2020).
4. Zheng, S. Q. *et al.* MotionCor2: anisotropic correction of beam-induced motion for improved cryo-electron microscopy. *Nature methods* **14,** 331–332; 10.1038/nmeth.4193 (2017).
5. Rohou, A. & Grigorieff, N. CTFFIND4: Fast and accurate defocus estimation from electron micrographs. *Journal of structural biology* **192,** 216–221; 10.1016/j.jsb.2015.08.008 (2015).
6. Wagner, T. & Raunser, S. The evolution of SPHIRE-crYOLO particle picking and its application in automated cryo-EM processing workflows. *Communications biology* **3,** 61; 10.1038/s42003-020-0790-y (2020).
7. Schöenfeld, F., Stabrin, M., Shaikh, T. R., Wagner, T. & Raunser, S. Accelerated 2D Classification With ISAC Using GPUs. *Frontiers in molecular biosciences* **9,** 919994; 10.3389/fmolb.2022.919994 (2022).
8. Yang, Z., Fang, J., Chittuluru, J., Asturias, F. J. & Penczek, P. A. Iterative stable alignment and clustering of 2D transmission electron microscope images. *Structure (London, England : 1993)* **20,** 237–247; 10.1016/j.str.2011.12.007. (2012).
9. Moriya, T. *et al.* High-resolution Single Particle Analysis from Electron Cryo-microscopy Images Using SPHIRE. *Journal of visualized experiments : JoVE*; 10.3791/55448. (2017).
10. Ciniawsky, S. *et al.* Molecular snapshots of the Pex1/6 AAA+ complex in action. *Nature communications* **6,** 7331; 10.1038/ncomms8331 (2015).
11. Scheres, S. H. W. RELION: implementation of a Bayesian approach to cryo-EM structure determination. *Journal of structural biology* **180,** 519–530; 10.1016/j.jsb.2012.09.006 (2012).
12. Zivanov, J. *et al.* New tools for automated high-resolution cryo-EM structure determination in RELION-3. *eLife* **7**; 10.7554/eLife.42166 (2018).
13. Adams, P. D. *et al.* PHENIX: a comprehensive Python-based system for macromolecular structure solution. *Acta crystallographica. Section D, Biological crystallography* **66,** 213–221; 10.1107/S0907444909052925 (2010).
14. Terwilliger, T. C., Sobolev, O. V., Afonine, P. V., Adams, P. D. & Read, R. J. Density modification of cryo-EM maps. *Acta crystallographica. Section D, Structural biology* **76,** 912–925; 10.1107/S205979832001061X (2020).
15. Tan, Y. Z. *et al.* Addressing preferred specimen orientation in single-particle cryo-EM through tilting. *Nature methods* **14,** 793–796; 10.1038/nmeth.4347 (2017).
16. Jumper, J. *et al.* Highly accurate protein structure prediction with AlphaFold. *Nature* **596,** 583–589; 10.1038/s41586-021-03819-2 (2021).
17. Pettersen, E. F. *et al.* UCSF ChimeraX: Structure visualization for researchers, educators, and developers. *Protein science : a publication of the Protein Society* **30,** 70–82; 10.1002/pro.3943 (2021).
18. Emsley, P. & Cowtan, K. Coot: model-building tools for molecular graphics. *Acta crystallographica. Section D, Biological crystallography* **60,** 2126–2132; 10.1107/S0907444904019158 (2004).
19. Pettersen, E. F. *et al.* UCSF Chimera--a visualization system for exploratory research and analysis. *Journal of computational chemistry* **25,** 1605–1612; 10.1002/jcc.20084 (2004).
20. Lopéz-Blanco, J. R. & Chacón, P. iMODFIT: efficient and robust flexible fitting based on vibrational analysis in internal coordinates. *Journal of structural biology* **184,** 261–270; 10.1016/j.jsb.2013.08.010 (2013).
21. Croll, T. I. ISOLDE: a physically realistic environment for model building into low-resolution electron-density maps. *Acta crystallographica. Section D, Structural biology* **74,** 519–530; 10.1107/S2059798318002425 (2018).

22. Krissinel, E. & Henrick, K. Inference of macromolecular assemblies from crystalline state. *Journal of molecular biology* **372,** 774–797; 10.1016/j.jmb.2007.05.022 (2007).
23. Laskowski, R. A. PDBsum: summaries and analyses of PDB structures. *Nucleic acids research* **29,** 221–222; 10.1093/nar/29.1.221 (2001).
24. Pan, M. *et al.* Mechanistic insight into substrate processing and allosteric inhibition of human p97. *Nature structural & molecular biology* **28,** 614–625; 10.1038/s41594-021-00617-2 (2021).
